# Remarkably high repeat content in the genomes of sparrows: the importance of genome assembly completeness for transposable element discovery

**DOI:** 10.1101/2023.10.26.564301

**Authors:** Phred M. Benham, Carla Cicero, Merly Escalona, Eric Beraut, Colin Fairbairn, Mohan P. A. Marimuthu, Oanh Nguyen, Ruta Sahasrabudhe, Benjamin L. King, W. Kelley Thomas, Adrienne I. Kovach, Michael W. Nachman, Rauri C. K. Bowie

**Author notes:** Corresponding author: Phred M. Benham.

## Abstract

Transposable elements (TE) play critical roles in shaping genome evolution. However, the highly repetitive sequence content of TEs is a major source of assembly gaps. This makes it difficult to decipher the impact of these elements on the dynamics of genome evolution. The increased capacity of long-read sequencing technologies to span highly repetitive regions of the genome should provide novel insights into patterns of TE diversity. Here we report the generation of highly contiguous reference genomes using PacBio long read and Omni-C technologies for three species of sparrows in the family Passerellidae. To assess the influence of sequencing technology on TE annotation, we compared these assemblies to three chromosome-level sparrow assemblies recently generated by the Vertebrate Genomes Project and nine other sparrow species generated using a variety of short- and long-read technologies. All long-read based assemblies were longer in length (range: 1.12-1.41 Gb) than short-read assemblies (0.91-1.08 Gb). Assembly length was strongly correlated with the amount of repeat content, with longer genomes showing much higher levels of repeat content than typically reported for the avian order Passeriformes. Repeat content for the Bell’s sparrow (31.2% of genome) was the highest level reported to date for a songbird genome assembly and was more in line with woodpecker (order Piciformes) genomes. CR1 LINE elements retained from an expansion that occurred 25-30 million years ago were the most abundant TEs in the song sparrow genome. Although the other five sparrow species also exhibit evidence for a spike in CR1 LINE activity at 25-30 million years ago, LTR elements stemming from more recent expansions were the most abundant elements in these species. LTRs were uniquely abundant in the Bell’s sparrow genome deriving from two recent peaks of activity. Higher levels of repeat content (79.2-93.7%) were found on the W chromosome relative to the Z (20.7-26.5) or autosomes (16.1-30.9%). These patterns support a dynamic model of transposable element expansion and contraction underpinning the seemingly constrained and small sized genomes of birds. Our work highlights how the resolution of difficult-to-assemble regions of the genome with new sequencing technologies promises to transform our understanding of avian genome evolution.

## INTRODUCTION

The dynamics of transposable element (TE) activity within host genomes are a major driver of genome evolution (Ågren & Wright 2011). Transposable elements proliferate throughout the genome either through copy-and-paste mechanisms (Class I elements; e.g. long interspersed nuclear elements [LINEs]) or through cut-and-paste mechanisms (class II elements; e.g. DNA transposons). The mobility of these elements in the genome contributes to structural variation (e.g. indels, inversions), alterations to gene expression, and the evolution of gene regulatory networks (Feschotte 2008; Schrader & Schmitz 2018). Given the genomic disruption potentially caused by TEs, the majority of new TE insertions are likely deleterious and species exhibit a wide range of defense mechanisms to both silence and delete TEs from the genome (Goodier 2016). Nevertheless, co-option of TEs by the host genome has led to the evolution of novel phenotypes (Mi et al. 2000; Cornelis et al. 2017) including color polymorphisms (van’t Hof et al. 2016; Kratochwil et al. 2022), increased immunity (Brosh et al. 2022), insecticide resistance (Daborn et al. 2002), and speciation (Serrato-Capuchina & Matute 2018). To date, much of TE biology has focused on model organisms with well-characterized genomic resources. The generation of high-quality genomes for a diversity of non-model organisms (Teeling et al. 2018; Feng et al. 2020; Rhie et al. 2021; Lewin et al. 2022) promises to broaden our understanding of how co-evolutionary dynamics between TEs and their host shape genome evolution.

Avian genomes provide an illuminating case of how expanding the diversity of available genome assemblies has altered our understanding of TE dynamics. Among amniotes, birds exhibit the smallest and most constrained genomes. Although contraction of avian genomes likely began prior to the evolution of flight (Organ et al. 2007), the high metabolic demand of flight is the leading hypothesis for continued constraint on avian genome size evolution (Hughes & Hughes 1995; Andrews et al. 2009; Wright et al. 2014). Consistent with a hypothesis of constrained genome evolution, the first avian genomes sequenced revealed low repeat content (<10%), little recent TE activity, and high chromosomal stability (Ellegren 2010). Detailed TE annotation of an increasing diversity of avian genome assemblies has since challenged the early narrative of low repeat content and high stability. First, comparative analyses across 12 avian genomes showed that the apparent stability in avian genome size was actually the product of a more dynamic history of genomic expansions offset by large-scale deletions (Kapusta et al. 2017). Second, extensive variation in the timing and proliferation of TE elements has been discovered across birds (Kapusta & Suh 2017; Suh et al. 2018; Galbraith et al. 2021). This includes the discovery of relatively high repeat content of 20-30% in the orders Piciformes (woodpeckers and allies) and Bucerotiformes (hornbills and hoopoes; Zhang et al. 2014; Manthey et al. 2018; Feng et al. 2020). Third, novel TEs have been discovered in avian lineages that derive from horizontal gene transfer from filarial nematodes (Suh et al. 2016). Finally, highly contiguous assemblies have confirmed that previous challenges to assembling the W chromosome were due in part to its role as a refugium for long terminal repeat (LTR) retrotransposons (Peona et al. 2021; Warmuth et al. 2022).

Our understanding of TE dynamics in avian genomes is poised to advance further with the increased use of long-read sequencing technologies (Kapusta & Suh 2017; Rhie et al. 2021). Repetitive regions of the genome, including centromeres, telomeres, and the W chromosome, are a major source of assembly gaps. Consequently, repetitive DNA is thought to make up a large proportion of the 7-42% of the genomic DNA missing from short-read genome assemblies relative to flow cytometry or densitometry estimates of genome size (hereafter the C-value; Peona et al. 2018). Indeed, a recent comparison of assembly methods for the Paradise Crow (*Lycocorax pyrrhopterus*) showed that gaps in short-read assemblies were primarily caused by LTR retrotransposons and simple repeats (Peona et al. 2020). Further, a recent comparison of activity levels of the chicken repeat 1 (CR1) retrotransposon across 117 avian genomes found a relationship between assembly contiguity (scaffold N50) and number of full length CR1s identified in individual genomes (Galbraith et al. 2021). This pattern was found across all genomes analyzed and also explained intra-generic variation in CR1 insertions. Detailed TE annotation of highly contiguous genomes will be essential for overcoming the confounding influence of assembly quality on patterns of TE diversity. In particular, studies leveraging highly contiguous genomes to explore TE dynamics across shorter evolutionary time scales are lacking but will be essential for understanding the contributions of these elements to the generation of avian diversity.

To this end, we performed in-depth TE annotations of highly contiguous genomes generated from six closely related sparrow species in the family Passerellidae. Passerellidae sparrows are a diverse clade of oscine Passeriformes, with 132 recognized species that are found throughout the Americas from northern Canada to southern Chile (Winkler et al. 2020). We generated *de novo* genome assemblies for Bell’s sparrow (*Artemisiospiza belli*), Savannah sparrow (*Passerculus sandwichensis*), and song sparrow (*Melospiza melodia*) for this paper as part of the California Conservation Genomics Project (CCGP; Shaffer et al. 2022). We analyze these assemblies alongside three genomes recently sequenced by the Vertebrate Genomes Project (VGP; Rhie et al. 2021) for saltmarsh (*Ammospiza caudacutus*), Nelson’s (*Ammospiza nelsoni*), and swamp sparrow (*Melospiza georgiana*), for a study of the genomic basis of tidal marsh adaptation. These new genome assemblies come from six members of the ‘grassland’ sparrow clade (Klicka et al. 2014). True to their name, all species can be generally found in a variety of shrub and grassland habitats across North America. Savannah and song sparrow are the two most ecologically and geographically widespread species occupying a broad range of tundra, alpine, meadow, prairie, marsh, and shrub habitats from Alaska and northern Canada south through Mexico to Guatemala (Arcese et al. 2002; Wheelwright & Rising 2008). Bell’s sparrow is found primarily in more arid chaparral and coastal sage habitat from northwestern California south into Baja California, Mexico and east into the southern San Joaquin valley and Mojave desert of southeastern California (Cicero & Koo 2012). Nelson’s and Swamp sparrow can primarily be found in central to eastern North America, principally in marsh habitats (Shriver et al. 2018; Herbert & Mowbray 2019). Saltmarsh sparrow is exclusively found in tidal marsh habitats of the Atlantic coast, and Nelson’s, swamp, song and Savannah sparrow all include tidal marsh specialist subspecies (Greenberg et al. 2006; Walsh et al. 2019a).

Song and Savannah sparrow are two of the most polytypic North America bird species, with 25 and 17 subspecies described in the song (Patten & Pruett 2009) and Savannah sparrow (Wheelwright & Rising 2008), respectively. In general, subspecific divergence across ecological gradients has long made all six species the focus of geographic variation and speciation studies (Marshall 1948; Aldrich 1984; Rising 2001; Cicero & Johnson 2006; Walsh et al. 2017; Mikles et al. 2020; Walsh et al. 2019b, 2021; Clark et al. 2022). The Bell’s sparrow also forms a narrow hybrid zone with the sagebrush sparrow (*Artemisiospiza nevadensis*) in the Owen’s Valley of eastern California (Cicero & Johnson 2007; Cicero & Koo 2012), while Nelson’s and saltmarsh sparrow hybridize along the coast of southern Maine (Rising & Avise 1993; Shriver et al. 2005; Walsh et al. 2015). Additionally, studies of these six sparrow species have provided important insights into avian life history and demography (Nice 1937; Johnston 1954; Keller et al. 1994; Keller & Arcese 1998; Marr et al. 2002; Freeman-Gallant et al. 2005; Ruskin et al. 2017a,b; Field et al. 2018), physiology (Poulson 1965; Greenberg et al. 2012; Benham & Cheviron 2020), vocal learning and behavior (Marler & Peters 1977; Searcy & Marler 1981; Williams et al. 2022), and migratory behavior (Moore et al. 1978; Able & Able 1996). The generation of highly contiguous reference genomes for these sparrow species with in-depth TE annotations will thus provide a critically important resource for future research in this intensively studied clade. In addition to the six new sparrow genome assemblies, nine other assemblies were analyzed from across the Passerellidae. The previous assemblies were produced using a variety of short- and long-read sequencing approaches. Previously sequenced genomes also include short-read assemblies for both the song and saltmarsh sparrows, which allows for intra-specific comparisons to assess the impact of sequencing technology on repeat annotation. We take advantage of the diverse genomic resources available from within this single avian family to ask: **(1)** what is the impact of sequencing technology and assembly completeness on TE element annotation? And **(2)** how do the evolutionary dynamics of TEs vary among closely related sparrow species. Addressing these questions will be important for determining how different sequencing approaches may introduce bias into comparative genomics analyses. In addition, our comparisons provide insights into how analyses based on short-read assemblies may miss important dynamics of avian genome evolution.

## METHODS

### CCGP genome sampling

We sequenced liver from an adult female Bell’s sparrow (*Artemisiospiza belli canescens*) collected on 25 June 2018 at Hunter Cabin, 1.5 Mi. east of Jackass Spring, Death Valley National Park, Inyo Co., California (36.54758°N, 117.48786°W; elevation: 6860 ft.). Blood and liver tissue samples were sequenced from an immature female song sparrow (*Melospiza melodia gouldii*) captured on 5 September 2020 in oak woodland habitat at Mitsui Ranch, Sonoma Mountain, Sonoma Co., California (38.33131°N; 122.57720°W). Liver tissue for sequencing was collected from an adult male Savannah sparrow (*Passerculus sandwichensis alaudinus*) on 20 May 2015 in tidal marsh habitat at the San Francisco Bay National Wildlife Refuge, Santa Clara Co., California (37° 26.029’N; 122° 0.996’W: elevation: 4 m). Bell’s and song sparrow individuals were collected with approval of California Department of Fish and Wildlife (CDFW permit #: SCP-458), the U.S. Fish and Wildlife Service (USFWS permit #: MB153526), Death Valley National Park (Bell’s sparrow only; permit#: DEVA-2015-SCI-0040), and following protocols approved by the University of California, Berkeley IACUC (AUP-2016-04-8665-1). The Savannah sparrow sample was also collected with approval from CDFW (permit #: SCP-012913), USFWS (permit #: MB24360B-0), the San Francisco Bay National Wildlife Refuge (special use permit: 2015-015), and using methods approved by the University of Illinois, Urbana-Champaign IACUC (protocol #: 13418). Voucher specimens were deposited at the Museum of Vertebrate Zoology, Berkeley, CA for the Bell’s sparrow (https://arctos.database.museum/guid/MVZ:Bird:192114) and song sparrow (https://arctos.database.museum/guid/MVZ:Bird:193390). A specimen voucher of the Savannah sparrow was deposited at the Field Museum of Natural History in Chicago, IL (FMNH:Birds:499929).

### DNA extraction, library preparation, and sequencing for CCGP genomes

High molecular weight (HMW) genomic DNA (gDNA) for PacBio HiFi library preparation was extracted from 27 mg and 15 mg of liver tissue from the Bell’s and Savannah sparrow samples, respectively. Extractions were performed using the Nanobind Tissue Big DNA kit following the manufacturer’s instructions (Pacific BioSciences - PacBio, Menlo Park, CA). For song sparrow, HMW gDNA was isolated from whole blood preserved in EDTA. A total of 30µl of whole blood was added to 2 ml of lysis buffer containing 100mM NaCl, 10 mM Tris-HCl pH 8.0, 25 mM EDTA, 0.5% (w/v) SDS and 100µg/ml Proteinase K. Lysis was carried out at room temperature for a few hours until the solution was homogenous. The lysate was treated with 20µg/ml RNase A at 37°C for 30 minutes and cleaned with equal volumes of phenol/chloroform using phase lock gels (Quantabio Cat # 2302830). DNA was precipitated by adding 0.4X volume of 5M ammonium acetate and 3X volume of ice-cold ethanol. The DNA pellet was washed twice with 70% ethanol and resuspended in an elution buffer (10mM Tris, pH 8.0), and purity was estimated using absorbance ratios (260/280 = 1.81-1.84 and 260/230 = 2.29-2.40) on a NanoDrop ND-1000 spectrophotometer. The final DNA yield (Bell’s: 13 μg; Savannah: 16 μg; song: 150 μg total) was quantified using the Quantus Fluorometer (QuantiFluor ONE dsDNA Dye assay; Promega, Madison, WI). The size distribution of the HMW DNA was estimated using the Femto Pulse system (Agilent, Santa Clara, CA): 62% of the fragments were >140 Kb for Bell’s sparrow; 60% of the fragments were >140 Kb for Savannah sparrow; and 85% of the DNA was found in fragments >120 Kb for song sparrow.

The HiFi SMRTbell library was constructed using the SMRTbell Express Template Prep Kit v2.0 following the manufacturer’s protocols (Pacific Biosciences - PacBio, Menlo Park, CA; Cat. #100-938-900). HMW gDNA was sheared to a target DNA size distribution between 15-20 Kb and concentrated using 0.45X of AMPure PB beads (PacBio Cat. #100-265-900) for the removal of single-strand overhangs at 37°C for 15 minutes. Enzymatic steps of DNA damage repair were performed at 37°C for 30 minutes, followed by the end repair and A-tailing steps at 20°C for 10 minutes and 65°C for 30 minutes. Ligation of overhang adapter v3 was performed at 20°C for 60 minutes with subsequent heating to 65°C for 10 minutes to inactivate the ligase. Finally, DNA product was nuclease treated at 37°C for 1 hour. To collect fragments greater than 9 Kb, the resulting SMRTbell library was purified and concentrated with 0.45X Ampure PB beads (PacBio, Cat. #100-265-900) for size selection using the BluePippin system (Sage Science, Beverly, MA; Cat #BLF7510). The 15-20 Kb average HiFi SMRTbell library was sequenced at the University of California Davis DNA Technologies Core (Davis, CA) using two 8M SMRT cells, Sequel II sequencing chemistry 2.0, and 30-hour movies each on a PacBio Sequel II sequencer.

The Omni-C library was prepared using the Dovetail^TM^ Omni-C^TM^ Kit (Dovetail Genomics, CA) according to the manufacturer’s protocol with slight modifications. First, specimen tissue was ground thoroughly with a mortar and pestle while cooled with liquid nitrogen. Subsequently, chromatin was fixed in place in the nucleus and then passed through 100 μm and 40 μm cell strainers to remove large debris. Fixed chromatin was digested under various conditions of DNase I until a suitable fragment length distribution of DNA molecules was obtained. Chromatin ends were repaired and ligated to a biotinylated bridge adapter followed by proximity ligation of adapter containing ends. After proximity ligation, crosslinks were reversed and the DNA purified from proteins. Purified DNA was treated to remove biotin that was not internal to ligated fragments. An NGS library was generated using an NEB Ultra II DNA Library Prep kit (NEB, Ipswich, MA) with an Illumina compatible y-adaptor. Biotin-containing fragments were then captured using streptavidin beads. The post-capture product was split into two replicates prior to PCR enrichment to preserve library complexity with each replicate receiving unique dual indices. The library was sequenced at the Vincent J. Coates Genomics Sequencing Lab (Berkeley, CA) on an Illumina NovaSeq platform (Illumina, San Diego, CA) to generate over 100 million 150 bp paired end reads per species. See supplemental Table S1 for details on PacBio and Illumina sequencing.

### Assembly of CCGP genomes

We assembled the genome of the three CCGP sparrows following the CCGP assembly pipeline Version 3.0 (see Lin et al. 2022; Supplemental Table S2). The pipeline takes advantage of long and highly accurate PacBio HiFi reads alongside chromatin capture Omni-C data to produce high quality and highly contiguous genome assemblies while minimizing manual curation.

In brief, we removed remnant adapter sequences from the PacBio HiFi dataset for all three assemblies using HiFiAdapterFilt (Sim et al. 2022) and obtained the initial dual assembly with the filtered PacBio reads using HiFiasm (Cheng et al. 2021). The dual assembly consists of a primary and alternate assembly: the primary assembly is more complete and consists of longer phased blocks, while the alternate consists of haplotigs (contigs with the same haplotype) in heterozygous regions and is more fragmented. Given the characteristics of the latter, it cannot be considered a complete assembly of its own but rather is a complement of the primary assembly (https://lh3.github.io/2021/04/17/concepts-in-phased-assemblies, https://www.ncbi.nlm.nih.gov/grc/help/definitions/).

Next, we identified sequences corresponding to haplotypic duplications, contig overlaps and repeats on the primary assembly with purge_dups (Guan et al 2020) and transferred these sequences to the corresponding alternate assembly. We aligned the Omni-C data to both assemblies following the Arima Genomics Mapping Pipeline (https://github.com/ArimaGenomics/mapping_pipeline) and used SALSA to produce scaffolds for the primary assembly (Ghurye et al. 2017, Ghurye et al. 2019).

Omni-C contact maps for the primary assembly were produced by aligning the Omni-C data with BWA-MEM (Li 2013), identifying ligation junctions, and generating Omni-C pairs using pairtools (Golobordko et al. 2018). We generated a multi-resolution Omni-C matrix with cooler (Abdennur an Mirny 2020) and balanced it with hicExplorer (Ramírez et al. 2018). To visualize and check contact maps for mis-assemblies, we used HiGlass (Kerpedjiev et al. 2018) and the PretextSuite (https://github.com/wtsi-hpag/PretextView; https://github.com/wtsi-hpag/PretextMap; https://github.com/wtsi-hpag/PretextSnapshot). In detail, if we identified a strong off-diagonal signal in the proximity of a join that was made by the scaffolder, and a lack of signal in the consecutive genomic region, we dissolved it by breaking the scaffolds at the coordinates of the join. After this process, no further manual joins were made. Some of the remaining gaps (joins generated by the scaffolder) were closed using the PacBio HiFi reads and YAGCloser (https://github.com/merlyescalona/yagcloser). We checked for contamination using the BlobToolKit Framework (Challis et al. 2020). Finally, upon submission of the assemblies to NCBI, we trimmed remnants of sequence adaptors and mitochondrial contamination identified during NCBI’s own contamination screening.

We assembled the mitochondrial genomes for each of the sparrows from their corresponding PacBio HiFi reads starting from the same mitochondrial sequence of *Zonotrichia albicollis* (NCBI:NC_053110.1; Feng et al. 2020, B10K Project Consortium) and using the reference-guided pipeline MitoHiFi (Uliano-Silva et al 2021; Allio et al. 2020). After completion of the nuclear genomes, we searched for matches of the resulting mitochondrial assembly sequence in their nuclear genome assembly using BLAST+ (Camacho et al. 2009), filtering out contigs and scaffolds from the nuclear genome with a sequence identity >99% and a size smaller than the mitochondrial assembly sequence.

### Genome assembly assessment

We generated k-mer counts from the PacBio HiFi reads using meryl (https://github.com/marbl/meryl). GenomeScope2.0 (Ranallo-Benavidez et al. 2020) was used to estimate genome features including genome size, heterozygosity, and repeat content from the resulting k-mer spectrum. To obtain general contiguity metrics, we ran QUAST (Gurevich et al. 2013). Genome quality and completeness were quantified with BUSCO (Manni et al. 2021) using the 8,338 genes in the Aves ortholog database (aves_odb10) for the three CCGP genomes. Assessment of base level accuracy (QV) and k-mer completeness was performed using the previously generated meryl database and merqury (Rhie et al. 2020). We further estimated genome assembly accuracy via a BUSCO gene set frameshift analysis using the pipeline described in Korlach et al. (2017). We followed the quality metric nomenclature established by Rhie et al. (2021), with the genome quality code x.y.Q.C, where, x = log10[contig NG50]; y = log10[scaffold NG50]; Q = Phred base accuracy QV (quality value); C = % genome represented by the first ‘n’ scaffolds, following a known karyotype of 2n=74 for *P. sandwichensis* (Bird Chromosome database - V3.0/2022; Degrandi et al 2020) and estimated 2n=80 for both *M. melodia* and *A. belli;* this estimation is the median number of chromosomes from closely related species (Genome on a Tree - GoaT; tax_tree(1729112); Challis et al. 2023). Quality metrics for the notation were calculated on the primary assemblies. Finally, we used the JupiterPlot pipeline (https://github.com/JustinChu/JupiterPlot) to visualize higher level synteny between the scaffolds of each sparrow assembly and chromosomes of the zebra finch (*Taeniopygia guttata*) genome assembly (Warren et al. 2010). Scaffolds representing 80% of each draft assembly (ng=80) were mapped to zebra finch chromosomes exceeding 1Mb in length (m=1000000).

### VGP genome sampling

The three CCGP were compared genomes to three chromosome-level assemblies of closely related sparrows generated by the Vertebrate Genomes Project following protocols outlined in Rhie et al. (2021). These samples were sequenced from blood samples (100 – 200 µl in ethanol) taken from female Nelson’s and swamp sparrows and a saltmarsh sparrow that was identified as a female in the field but lacked the W chromosome in the final assembly. The Nelson’s sparrow sample was collected by Nicole Guido (Saltmarsh Habitat and Avian Research Program) in South Branch Marsh River, Waldo County, Maine (44.5864°N; 68.8591°W) on 31 July 2020. The swamp sparrow sample was collected by Jonathan Clark (University of New Hampshire) in Durham, Rockingham County, New Hampshire (43.14°N; 71.00°W) on 23 July 2020. The saltmarsh sparrow sample was collected by Chris Elphick (University of Connecticut) from Barn Island Wildlife Management Area, New London County, Connecticut (41.338°N; 71.8677°W) on August 19 2020. Sample collection occurred under permits of the Maine Division of Inland Fisheries and Wildlife (#2020-314), Connecticut Department of Energy and Environmental Protection (#0221012b), and New Hampshire Fish and Game, and followed protocols approved under the University of New Hampshire IACUC (#190401). All individuals were released at the capture location immediately after sampling. DNA extracted from these three species was sequenced to 31.6-34.6x coverage of PacBio Sequel II HiFi long reads, 254-450x coverage of Bionano Genomics DLS, and 103-112x coverage for Arima Hi-C v2. Genome assemblies were generated from these data with the VGP standard assembly pipeline version 2.0, which included hifiasm v0.15.4, purge_dups v. 1.2.5, solve v. 3.6.1, salsa v. 2.3. Further details on raw data, sequence evaluations and curated assemblies can be found on VGP Genome Ark pages for the Nelson’s sparrow (https://genomeark.github.io/genomeark-all/Ammospiza_nelsoni.html), saltmarsh sparrow (https://genomeark.github.io/genomeark-all/Ammodramus_caudacutus.html), and swamp sparrow (https://genomeark.github.io/genomeark-all/Melospiza_georgiana.html).

### Genome size variation within sparrows

We estimated the distribution of genome sizes in Passerellidae sparrows with c-value data from 33 individuals of 21 species archived in the Animal Genome Size Database (Gregory 2022). The majority of these C-value estimates were generated using Feulgen image analysis densitometry with original values reported in Andrews et al. (2009) and Wright et al. (2014). This includes all species for which we compared C-values to assembly lengths. C-value estimates of genome size were converted from picograms to base pairs using a conversion of 1pg = 0.978 Gb (Dolezel et al. 2003). We additionally obtained genome assembly length from nine publicly available assemblies that were sequenced previously for members of the Passerellidae. Six of these were based on Illumina short-read sequence data and include a second song sparrow genome from Alaska (Louha et al. 2019), a short-read genome of the saltmarsh sparrow (*Ammospiza caudacuta*; Walsh et al. 2019a), plus genomes for the white-throated sparrow (*Zonotrichia albicollis*; Tuttle et al. 2016), dark-eyed junco (*Junco hyemalis*; Friis et al. 2022), chipping sparrow (*Spizella passerina*; Feng et al. 2020), and grasshopper sparrow (*Ammodramus savannarum;* Carneiro 2021). The remaining three assemblies were generated using PacBio long-read sequence data and include a third song sparrow genome from British Columbia (Feng et al. 2020), a white-crowned sparrow genome (*Zonotrichia leucophrys*; unpublished), and a contig-level assembly of the California towhee (*Melozone crissalis;* Black et al. 2023). For complete GenBank accession details, sequencing, and assembly methods for these genomes see supplemental Table S3. We compared C-value estimates of genome size and genome assembly length for all of the assemblies that included both these estimates of genome size (10 of 15 total).

### Repeat annotation

We performed detailed *de novo* repeat annotation and manual curation of repeat libraries for the Bell’s, song, and Savannah sparrow genomes sequenced by the CCGP in the program RepeatModeler2 with the ltrstruct option selected to improve identification of LTR elements (Flynn et al. 2020). Consensus transposable element libraries generated from RepeatModeler2 were then curated manually following protocols and methods of Goubert et al. (2022). First, we removed any redundancy in the *de novo* repeat libraries using cd-hit-est (Li & Godzik 2006) to cluster any consensus sequences together that were 80 base pairs in length and shared >80% similarity over more than 80% of their length. This corresponds to the 80-80-80 rule of Wicker et al. (2007) frequently used to classify TE elements as a single family. We then prioritized elements for manual curation that were at least 1000 base pairs in length and had at least 10 blastn hits in the genome assembly. For each consensus sequence prioritized for manual curation, we used blastn (Camacho et al. 2009) to identify other members of each TE family in the genome, and for each blastn hit we added 2000 bp of flanking sequence to both ends of the sequence. We then aligned the extended sequences using mafft (Katoh & Standley 2013) and removed gaps automatically using T-coffee (Notredame et al. 2000). The multiple sequence alignment produced by mafft was visualized in aliview (Larsson et al. 2014), and the termini of each element were identified based on canonical motifs of different element classes (e.g. 5’ TG and 3’ CA dinucleotides in LTR elements). A consensus sequence of the trimmed multiple sequence alignment was then generated using the cons tool in EMBOSS (Rice et al. 2000). Finally, the program TE-Aid (https://github.com/clemgoub/TE-Aid) was used to confirm structural properties and the presence of open reading frames for the expected proteins characteristic of each class of TE element. This process of blast, extension, and alignment was repeated iteratively for each element until termini were discovered. Following manual curation of TE sequences, we again used cd-hit-est with the same settings as above to cluster sequences belonging to the same family. The final set of manually curated sequences was then compared against a library of avian TE elements downloaded from repbase as well as other recently published TE datasets (e.g. *Dromaius novaehollandiae*, Peona et al. 2020) using cd-hit-est and the 80-80-80 rule above as a threshold to classify curated elements as belonging to previously identified TE families. We assigned the following species-specific prefixes for newly identified repeat elements in each sparrow species: pasSan (*Passerculus sandwichensis*), melMel (*Melospiza melodia*), and artBel (*Artemisiospiza belli*). For new TE families shared across two or more of the sparrow species we assigned the prefix Passerellidae. For each element the prefix was followed by the superfamily identity (e.g., LTR/ERV1). For elements where we could not confidently identify the complete consensus sequence we added the suffix .inc.

Curated TE libraries for all three sparrow species were merged into a single Passerellidae repeat library that was then used to annotate transposable element diversity in the genome assemblies of the three CCGP sparrow genomes using RepeatMasker v. 4.1.2 (Smit et al. 2015). We additionally used the *de novo* Passerellidae TE library to annotate the genome assemblies of the 3 VGP sparrow genomes and the 9 previously sequenced sparrow genomes available on Genbank. Secondly, for seven sparrow species with a contig N50 >1Mb and at least a scaffold level assembly, we performed separate RepeatMasker runs on the autosomes, Z chromosome, and W chromosome (if present). For the CCGP genomes, scaffolds were assigned to these different chromosomes based on homology with the Zebra Finch chromosomes using Minimap2 (Li 2018). Finally, we assessed temporal patterns of TE activity for these seven sparrow species. For autosomal, Z, and W chromosomes, we used the calcDivergenceFromAlign.pl script in the RepeatMasker package to estimate the Kimura 2-parameter (K2P) distance of each TE element from the consensus sequence. K2P distances were used to generate barplots for the LTR, SINE, LINE, and DNA classes of TE elements found in the genome. For autosomal loci, we additionally used a mutation rate estimate of 2.3×10^-9^ per generation from another Passerine species (*Ficedula albicollis,* Smeds et al. 2016) to estimate the approximate timing of repeat activity.

### Sparrow phylogeny construction

To further contextualize the history of transposable element proliferation across sparrows, we constructed a time-calibrated phylogeny using ultra-conserved elements (UCE) extracted from all available sparrow genomes (14 taxa including 12 species and 3 song sparrow subspecies) as well as the medium ground finch (*Geospiza fortis*), island canary (*Serinus canaria*), and zebra finch (*Taeniopygia guttata*) as outgroup taxa. We used the UCE-5k-probe-set and the phyluce pipeline (Faircloth 2016) to align probes to each genome, extended sequences by 1000 bp on either flank, aligned sequences using mafft v. 7.49 (Katoh & Standley 2013), and produced a final PHYLIP file of the UCEs. We used RAxML-NG (Stamatakis 2014) on the CIPRES science portal (Miller et al. 2010) to generate a maximum likelihood phylogeny of the concatenated UCE loci using the GTRCAT model, rapid bootstrapping, and the autoMRE bootstrapping criterion. Secondly, we used MCMCtree to estimate a time-calibrated phylogeny based on the topology generated by RAxML-NG. We used the fossil sparrow *Ammodramus hatcheri* (Steadman 1981) to calibrate the node between the grasshopper sparrow (*Ammodramus savannarum*) and all other sparrows. Following Oliveros et al. (2019), we set the calibration time to 12 Mya bounded by a minimum age of 7.5 Mya and a maximum age of 18.6 Mya. We implemented a model of independent rates among branches and drawing from a log-normal distribution. We performed two independent runs of MCMCtree with each run starting from a different random seed. The first 50,000 iterations were removed as burnin before running for another 100 million iterations with a sample taken every 1000 iterations. We assessed convergence of the two runs using tracer v. 1.6.0 (Rambaut et al. 2018).

## RESULTS

### Genome assemblies

The three CCGP assemblies included a low of 337 scaffolds in the Savannah sparrow and a high of 1,339 scaffolds in Bell’s sparrow; total assembly lengths ranged from 1.15-1.40 Gb (Table 1; Supplemental figures S1-S3). All assembly metrics indicate that the genomes are highly contiguous with contig N50 ranging from 5.98-8.31 Mb and scaffold N50 from 17.08-25.78 Mb. The largest contig length was over 32.13 Mb and the longest scaffold over 99.81 Mb. Over 96% of the genes in the avian orthologous database were found to be complete and single copy in BUSCO. These metrics indicate that the genomes generated *de novo* by the CCGP pipeline are in line with overall contiguity and completeness metrics for the three genomes generated by the VGP. Genomes for swamp, saltmarsh, and Nelson’s sparrow showed similar contig N50 ranging from 8.25-12.04 Mb, but scaffold N50s were approximately 3x as large (74.25-78.44 Mb). The VGP genomes were also assembled into chromosome-level assemblies with 36-40 chromosomes identified based on decreasing order of size. Finally, BUSCO scores were slightly higher for the VGP genomes, exceeding 98% complete and single copy genes in all three species, but this could also reflect the different databases used in BUSCO analyses of the CCGP and VGP genomes. Jupiter plots showed CCGP sparrow scaffolds mapping to most chromosomes of the zebra finch genome, with little evidence for inversions or translocations that may be indicative of misassemblies (Fig. 1). Similarly, although contact maps for the primary assemblies of the three CCGP genomes show some level of fragmentation, they also reveal little evidence for inversions or translocations. (Supplementary Figure S4). Given their greater contiguity, we only describe the primary assemblies here, but the sequences corresponding to both primary and alternate assemblies for each of the CCGP species are available on NCBI (See supplemental Table S4 and Data availability for details).

**Figure 1:**
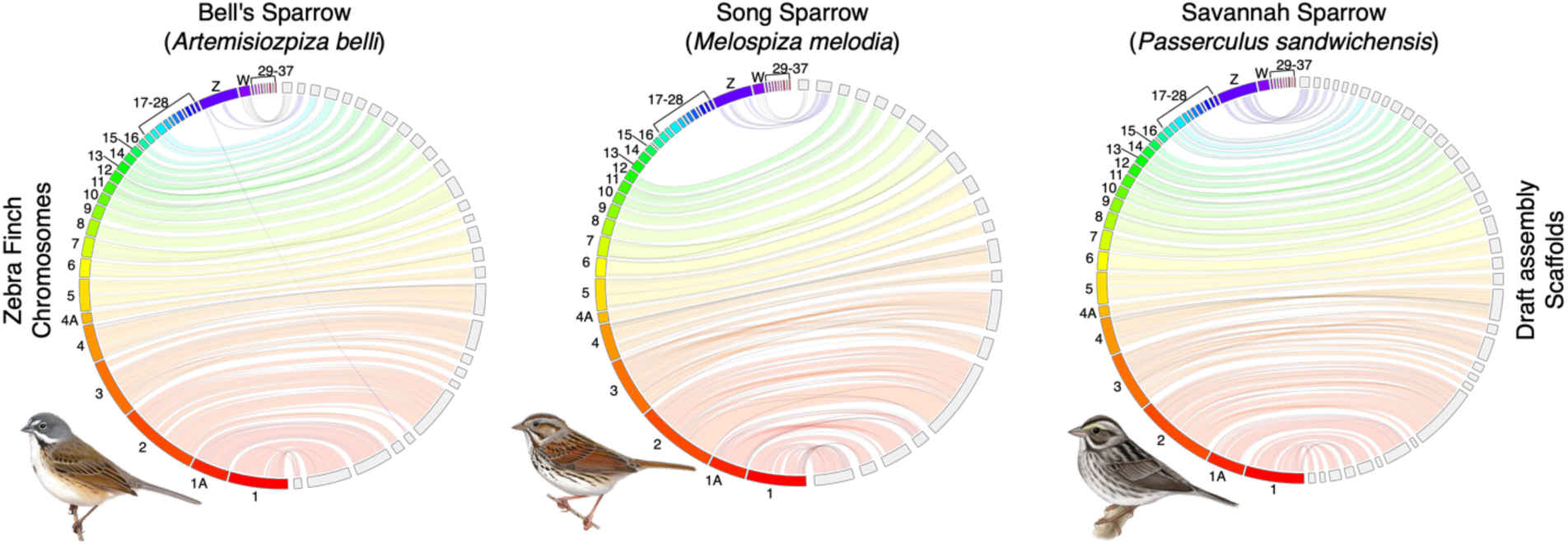
Jupiter plot comparing higher level synteny and completeness between the zebra finch (*Taeniopygia guttata*) genome (bTaeGut.4) and each of the three CCGP draft assemblies of Passerellidae sparrow species. Zebra finch chromosomes are on the left in each plot (colored) and sparrow scaffolds are on the right (light gray). Twists represent reversed orientation of scaffolds between assemblies. Song and Bell’s sparrow reference genome samples were both from females, whereas the Savannah sparrow reference was from a male. Song and Bell’s sparrow illustrations reproduced with the permission of https://birdsoftheworld.org with permission from Lynx Edicions. Savannah sparrow illustration contributed by Jillian Nichol Ditner.

**Table 1:**
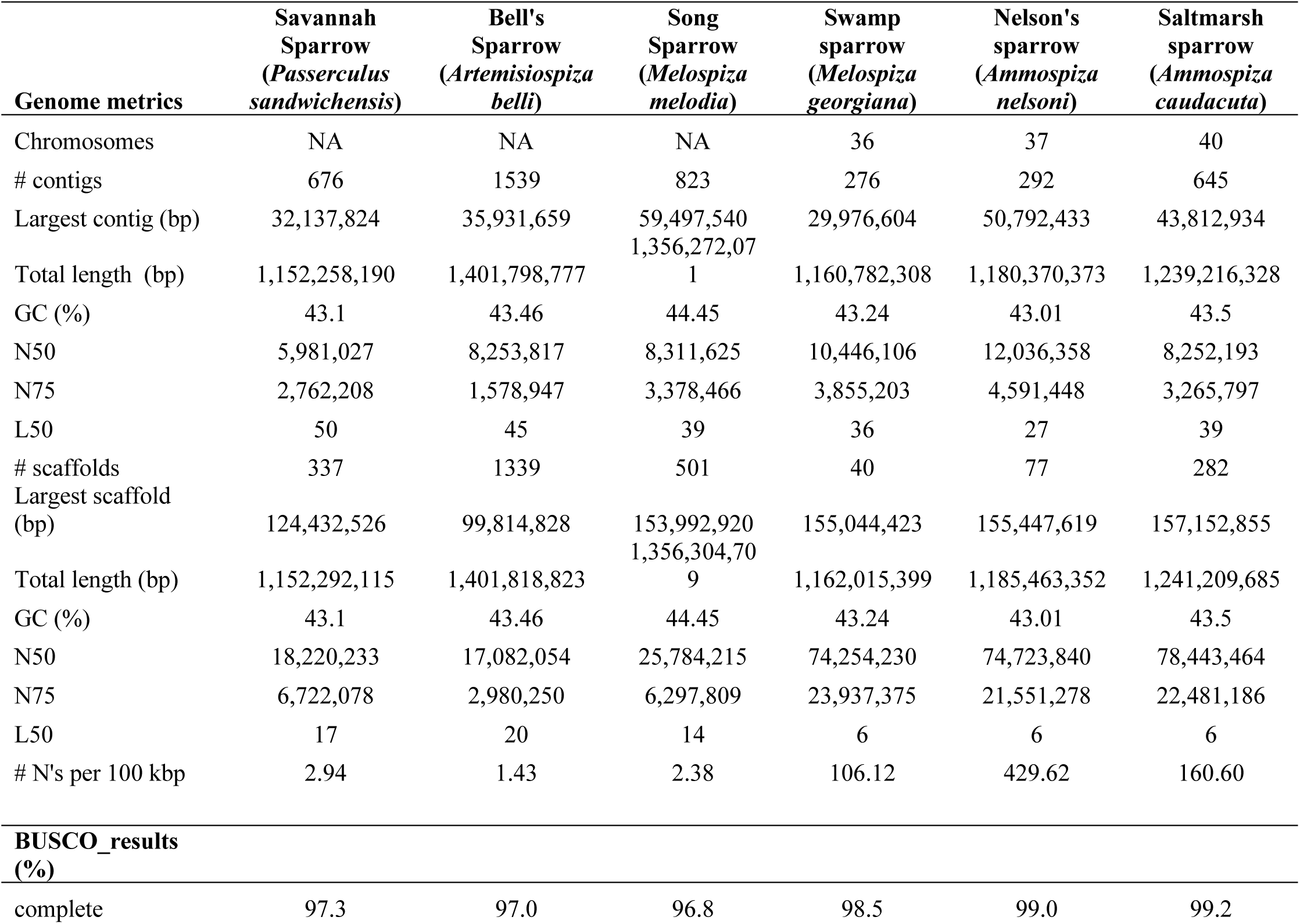
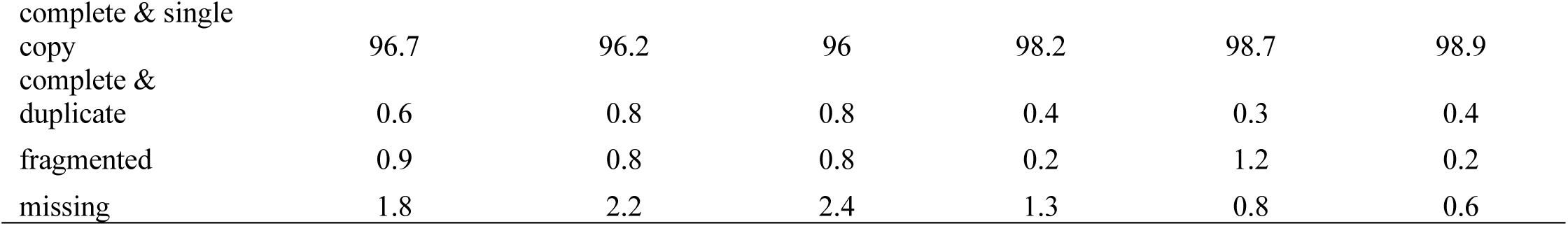
Comparison of assembly quality statistics and BUSCO search results among the three CCGP (left three) and three VGP (right three) genomes. BUSCO results for the CCGP genomes were obtained using the 8,338 universal single copy genes in birds found in the aves_odb10 database. BUSCO results from the VGP genomes were obtained using the 10,844 genes from the passeriformes_odb10 database.

### Genome size variation in sparrows

Adjusted genome size estimates from C-values of Passerellidae sparrows varied by 0.5 Gb from 1.13 Gb in Savannah sparrow (*Passerculus sandwichensis*) to 1.63 Gb in gray-browed brushfinch (*Arremon assimilis*), with a mean of 1.36 Gb (Fig. 2a). Previous short-read genome assemblies of sparrows varied in length from 0.91 Gb in the grasshopper sparrow to 1.05 Gb in the white-throated sparrow assembly, which were 0.16 to 0.42 Gb smaller than the corresponding C-value estimates of genome size for these species. Assembly lengths for the CCGP and VGP sparrow genomes varied from 1.15 Gb in the Savannah sparrow to 1.40 Gb in the Bell’s sparrow (Fig. 2a). Recently released long-read assemblies of the white-crowned sparrow (1.12 Gb) and California towhee (1.41 Gb) span a similar range. The length of the assemblies reported here closely approximated the C-value estimates of genome size for the Savannah sparrow (mean 1.18 Gb vs. 1.15 Gb assembly) and the song sparrow (mean 1.40 Gb vs. 1.35 Gb assembly), but was more divergent in the swamp sparrow (1.46 Gb vs. 1.16 Gb assembly). No C-value estimates exist for the other three sparrow species; however, alternate estimates of genome size are available from the kmer profiles analyzed in GenomeScope (supplemental Fig. S1-S3; https://www.genomeark.org/genomeark-all/). These profiles suggest that the Nelson’s (assembly: 1.18 Gb; GenomeScope: 1.19 Gb) and saltmarsh sparrow (1.24 vs. 1.22) assemblies closely match the expected genome length estimated from the kmer profile. In contrast, all three CCGP sparrow genome assemblies exceed the kmer profile genome size estimates (for example Bell’s sparrow assembly: 1.40 Gb vs. GenomeScope estimate: 1.13); whereas, the curated swamp sparrow assembly (1.16 Gb) was considerably shorter than the estimated length from GenomeScope (1.33 Gb). Together these data underscore the high level of completeness of the assemblies generated using long-read approaches, with less than 3% of the genome missing from most species. The swamp sparrow assembly appears to be an exception with 12-20% of the genome content potentially missing (depending on GenomeScope or C-value estimate). Purged repeat content may explain some of the missing data from the swamp sparrow assembly. Repeat content in the swamp sparrow was estimated to span 18.26% of the genome using our Passerellidae repeat library in RepeatMasker, while k-mer estimates of repeat content were 30.4% from GenomeScope. In contrast, prior sparrow genome assemblies that used primarily short-read data were inferred to be missing as much as 12-30% of the genomic DNA (Fig. 2b).

**Figure 2:**
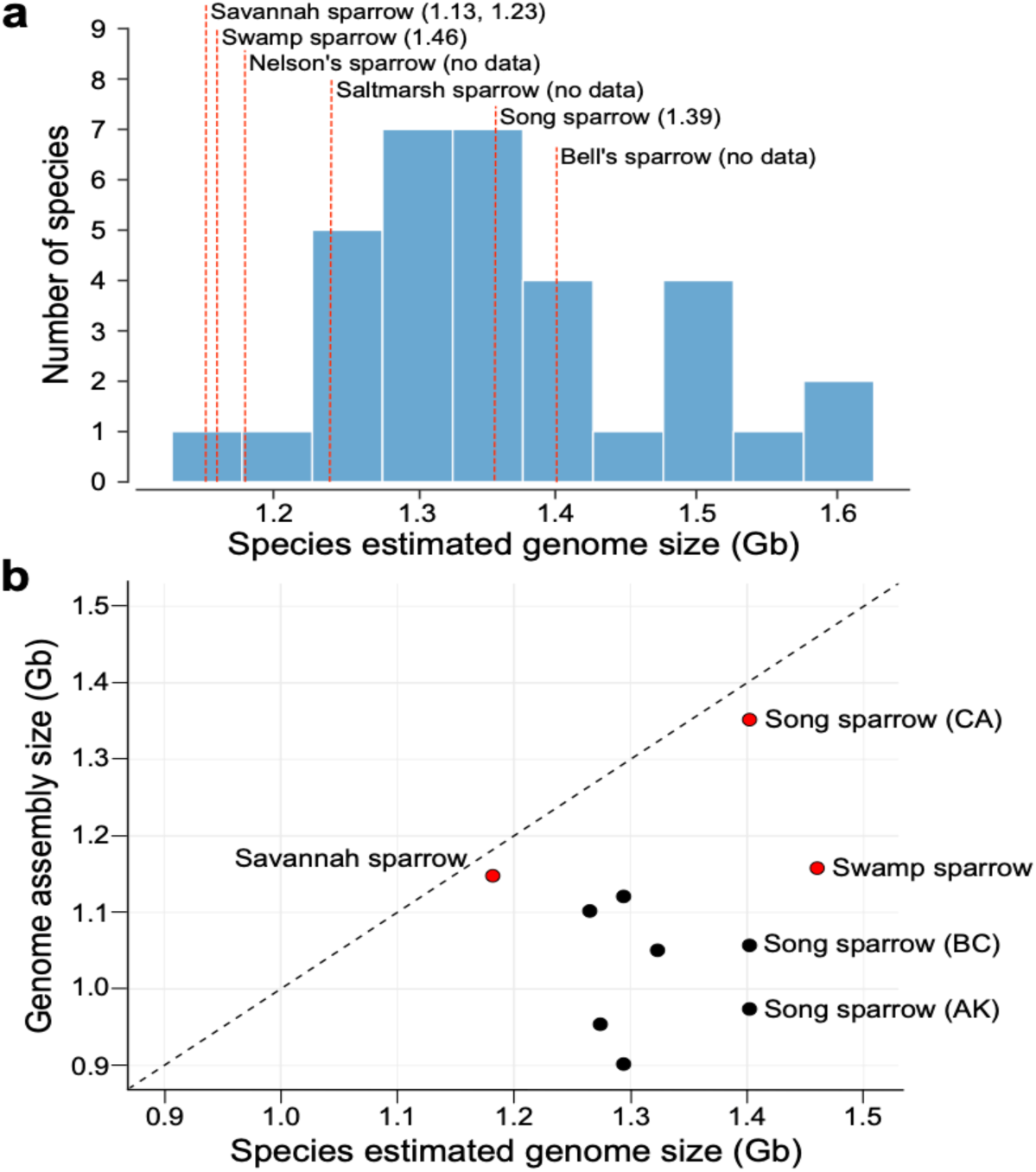
(a) Histogram showing variation in the genome sizes for 21 species (33 individuals) of Passerellidae sparrows estimated from Feulgen image analysis densitometry methods. C-values were adjusted based on the assumption that 1pg = 0.978 Gb (Dolezel et al. 2003). Genome size of the draft assemblies generated through the CCGP and VGP shown as red dashed lines with corrected c-value estimate (Gb) in parentheses. For *Passerculus sandwichensis* two estimates of genome size were found in the Animal Genome Size Database. (**b**) Comparison of estimated genome size from C-values versus genome assembly size (Gb). Black dashed line indicates the 1-to-1 line indicating equal estimates of genome size from the two metrics. Red dots are newly generated assemblies reported here. Nelson’s, saltmarsh, and Bell’s sparrow lack independent estimates of C-values and are not shown on this plot. Black dots denote previously published genome assemblies, which all show a shorter assembly length relative to the C-value estimate of genome size.

### Passerellidae de novo repeat library

*De novo* identification of transposable elements in RepeatModeler2 followed by manual curation led to the identification of 514, 704, and 650 TE families within the Savannah, song, and Bell’s sparrows, respectively. Merging of the three sparrow libraries produced a final Passerellidae TE library with 1,272 elements. This includes 361 elements shared by two or more sparrow species and 234, 341, and 336 elements unique to Savannah, song, and Bell’s sparrows, respectively. Similar to other avian species, LINE and LTR elements represent the majority of TEs identified in these sparrow species. These include 15 shared LINE families and 68 shared LTR families across all three species. Song sparrows had the most unique LINE elements (n=58), whereas Bell’s sparrow had the most unique LTR elements (n=122). Savannah sparrow had the least number of both elements identified (supplemental Fig. S5).

The curated Passerellidae repeat library was used to annotate all 15 sparrow assemblies. Annotation results from RepeatMasker showed that repeat content comprises a considerably higher percentage of the genome in the more contiguous, long-read assemblies (Fig. 3a). Bell’s sparrow, song sparrow, and California towhee showed the greatest proportion of repeat content with repeats comprising over 29% of the genome (Table 2; Fig. 3a). The Savannah sparrow exhibited the lowest proportion of repeats (16.5%) among the long-read assemblies, but this still exceeded the 6.5-10.3% of the genome covered by repeat content identified in other sparrow species. Indeed, we found contig N50 to be highly predictive of total repeat content discovered in genome assemblies. All eight sparrow assemblies with a contig N50 greater than 1 Mb had significantly more repeat content than assemblies with contig N50 less than 1 Mb (Fig. 3b; t=-6.174; df=7.8; p=0.0003). All assemblies with a contig N50 > 1 Mb were assembled using PacBio long-reads; whereas only 1 of 7 of the assemblies with contig N50 less than 1 Mb included PacBio long-read sequencing in the assembly. Variation in assembly length among species also strongly predicted repeat content (adjusted R^2^=0.95, p-value <<0.0001; Fig. 4c). This pattern was replicated among different assemblies of the song sparrow and saltmarsh sparrow. Repeat content increased from 7.1% (0.978 Gb assembly) to 10.3% (1.06 Gb assembly) to 29.5% (1.36 Gb assembly) as assembly length increased in the three song sparrow assemblies. In the saltmarsh sparrow, repeat content more than doubled from 10.6% in the short-read assembly (1.07 Gb) to 24.2% in the long-read assembly (1.24 Gb). Finally, the amount of repeat content significantly decreased as the percent missing DNA increased for each assembly with missing DNA inferred from the difference between C-value estimate and assembly length (adjusted R^2^=0.57, p-value = 0.007; Fig. 3d).

**Figure 3:**
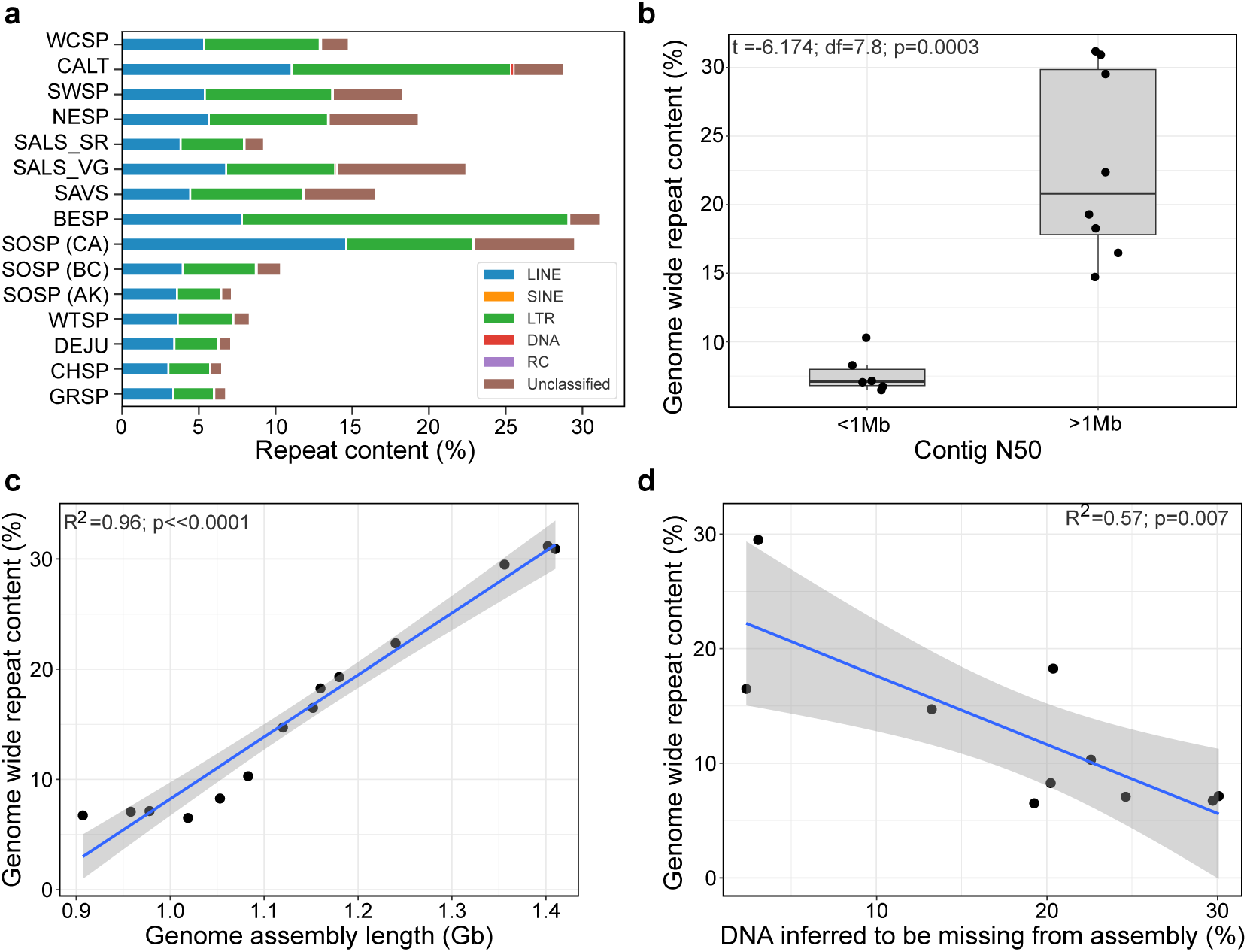
(a) Percentage of the genome comprising interspersed repeats, including: retroelements (LINE, SINE, LTR), DNA transposons (DNA), rolling-circles (RC), and unclassified elements. (white-crowned sparrow: WCSP; California towhee: CALT; swamp sparrow: SWSP; Nelson’s sparrow: NESP; saltmarsh sparrow short-read: SALS_SR; saltmarsh sparrow long-read: SALS_VG; Savannah sparrow: SAVS; Bell’s sparrow: BESP; song sparrow: SOSP; white-throated sparrow: WTSP; dark-eyed junco: DEJU; chipping sparrow: CHSP; grasshopper sparrow: GRSP). (**b**) The relationship between contig N50 and genome wide repeat content. Significantly higher levels of repeat content were discovered in genomes with a contig N50 greater than 1 Mb. All of which were generated with PacBio long-read technology. **(c**) Correlation between percent repeat content identified in each genome and the length of the assembled genome in Gb. (**d**) Correlation between percent repeat content and the amount of DNA inferred to be missing from each of the sparrow assemblies. C-value is assumed to be the more accurate estimate of total genome length. Percent missing DNA from each sparrow assembly is estimated as the difference between the c-value and assembly length.

**Figure 4:**
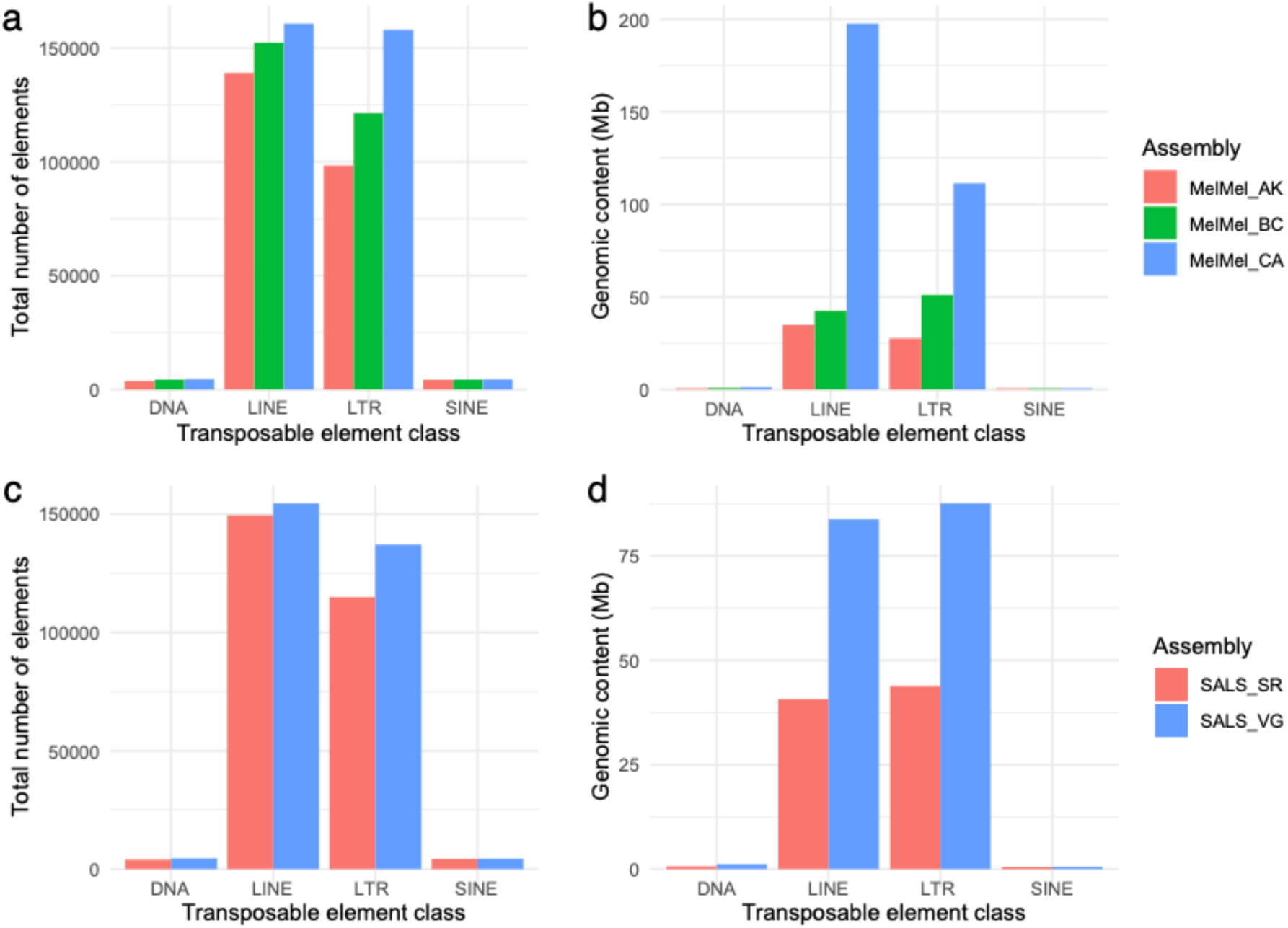
(**a-b**) Comparison of transposable element annotation across three song sparrow (*Melospiza melodia*) assemblies. (**a**) Total number of transposable elements found in each assembly. (**b**) Total genomic content (Mb) of transposable element content identified in each of the three song sparrow assemblies. MelMel_AK (red) is an assembly from an Alaskan bird sequenced using short-read and Chicago library technology. MelMel_BC (green) is a bird from British Columbia sequenced using Illumina short-read and PacBio SMRT long-read approaches MelMel_CA was generated using Hi-C and PacBio long read approaches for this paper. (**c-d)** Comparison of TE annotations in two saltmarsh sparrow assemblies (*Ammospiza caudacuta*). (**c**) Total number of transposable elements found in each assembly. (**b**) Total genomic content (Mb) of transposable element content identified in each of the three song sparrow assemblies. SALS_SR (Red) was assembled from Illumina short reads and the SALS_VG (blue) assembly was assembled using PacBio long read and Omni-C approaches with the Vertebrate Genomics Project pipeline.

**Table 2:** Percentage of each genome spanned by different classes of repeats. Estimates of each class of repeat region identified within RepeatMasker using the sparrow TE libraries generated *de novo* with RepeatModeler2.

| Element class: | Savannah<br>sparrow<br>(*Passerculus*<br>*sandwichensis*) | Bell's sparrow<br>(*Artemisiospiza*<br>*belli*) | Song<br>sparrow<br>(*Melospiza*<br>*melodia*) | Swamp<br>sparrow<br>(*Melospiza*<br>*georgiana*) | Nelson's<br>sparrow<br>(*Ammospiza*<br>*nelsoni*) | Saltmarsh<br>sparrow<br>(*Ammospiza*<br>*caudacuta*) |
| --- | --- | --- | --- | --- | --- | --- |
| LINE | 4.41 | 7.8 | 14.58 | 5.37 | 5.63 | 6.75 |
| SINE | 0.05 | 0.04 | 0.02 | 0.05 | 0.05 | 0.05 |
| LTR | 7.28 | 21.21 | 8.22 | 8.25 | 7.72 | 7.06 |
| DNA transposons | 0.09 | 0.10 | 0.09 | 0.08 | 0.08 | 0.1 |
| Rolling-circles | 0.02 | 0.02 | 0.01 | 0.01 | 0.02 | 0.05 |
| Unclassified | 4.66 | 2.01 | 6.56 | 4.51 | 5.81 | 8.40 |
| **Total**<br>**interspersed**<br>**repeats:** | 16.49 | 31.16 | 29.49 | 18.26 | 19.29 | 22.35 |
| Autosomal<br>chromosomes: | 16.17 | 30.91 | 28.84 | 15.85 | 16.11 | 19.48 |
| Z chromosome: | 20.72 | 23.75 | 25.28 | 22.40 | 26.42 | 26.46 |
| W chromosome: | NA | 82.59 | 91.03 | 93.73 | 79.23 | NA |
| **Other repeat**<br>**regions:** |  |  |  |  |  |  |
| Small RNA | 0.04 | 0.04 | 0.03 | 0.06 | 0.05 | 0.04 |
| Satellites | 0.33 | 0.21 | 0.24 | 0.25 | 0.22 | 0.25 |
| Simple repeats | 1.35 | 1.11 | 1.05 | 1.25 | 1.19 | 1.21 |
| Low complexity | 0.25 | 0.20 | 0.20 | 0.23 | 0.28 | 0.32 |

The comparison among the three song sparrow genome assemblies showed that LINEs and LTRs comprised the greatest number and total base pairs of newly discovered TE sequence (Fig. 4a). We found an additional 8,375 (a ∼5% increase) line elements in the California versus British Columbia song sparrow assemblies. Despite only a small increase in the total number of elements, we find that these LINE elements span an additional 155 Mb of DNA in the California song sparrow genome (Fig. 4b). This discrepancy likely stems both from different elements segregating at different frequencies in each population and LINEs in the California genome being of greater length on average. One of the most abundant LINE elements in the California song sparrow genome was found across all three of the CCGP sparrow genomes but was missing from the British Columbia song sparrow genome. This element was >6500 bp in length and nearly 4000 full length copies were found across the California song sparrow genome. Comparisons between a short-read and long-read assembly of the saltmarsh sparrow revealed a similar pattern (Fig. 4c-d). A small increase in the total number of LINE (∼3%) and LTR (∼16%) elements led to respective increases of 43.1 Mb and 43.8 Mb of TE DNA discovered in these genomes. These results further support the inference that missing DNA from previous assemblies corresponds to longer TE elements and may have been a major contributor to gaps.

The prevalence of different TE classes varied across our three genome assemblies. LTRs were the most abundant element within all sparrow genomes except the song sparrow assembly where LINE elements were the most abundant (14.58%; Table 2). Across different chromosomes, the density of repeat content (79.23-93.73%) was highest on the W chromosomes. W chromosome repeat content was particularly high in the genus *Melospiza* with over 90% of the W chromosome spanned by repetitive elements in both the song and swamp sparrow. Z chromosome repeat content tended to be higher than autosomal repeat content for all species except song and Bell’s sparrow (Table 2).

### Timing of repeat proliferation

We extracted an average of 4,663 (range: 3,699-4,839) ultra-conserved element (UCE) loci from the 17 reference genomes queried (Supplemental Table S5). From these loci we constructed a concatenated data matrix of 4,196 UCE loci shared across 95% of samples with a total of 4,815,326 base pairs. The concatenated maximum likelihood tree was well-resolved with all nodes receiving bootstrap support of 100. This topology was used as input into MCMCtree to estimate divergence times among the focal species (see supplemental Figure S6 for full phylogeny). This time-calibrated phylogeny indicated that the white-crowned sparrow split from the other sequenced sparrow species at 13.3 Mya (95% HPD: 7.4-17.8; Fig. 5), Bell’s sparrow diverged from other species in the grassland sparrow clade 7.9 Mya (95% HPD: 4.5-10.8), *Ammospiza* sparrows (Nelson’s and saltmarsh) split from Savannah, swamp, and song sparrow 6.8 Mya (95% HPD: 3.8-9.3), and Savannah sparrow diverged from the *Melospiza* sparrows 5.8 Mya (95% HPD: 3.3-7.9). Within the context of these divergence times, members of the Passerellidae family show sharply divergent histories of transposable element proliferation (Fig. 5). All members of the grassland sparrow clade show evidence for a spike in LINE element activity in the autosomes ∼25-30 Ma. In contrast, the white-crowned sparrow does not show evidence for this spike, but rather shows a normal distribution of LINE element divergence centered at ∼40-50 Ma. Although the timing of LINE proliferation in grassland sparrows appears to predate divergence estimates among Passerellidae species (Fig. 5), the contrast with white-crowned sparrow suggests it may have occurred more recently following divergence of these different sparrow lineages. Despite shared evidence for this period of LINE activity, song sparrows have retained more LINE elements from this proliferation (∼14% of the genome) than the other five sparrow species (only 1-3% of genomes). Bell’s sparrow shows a unique pulse of LTR proliferation approximately 12 Mya and a very recent (<5 Mya) proliferation of both LINE and LTR elements in the autosomes. For species with an assembled W chromosome, all show a steady accumulation of LTR elements on the W chromosome, with endogenous retroviruses being the most prolific and representing up to a maximum of 69.2% in the song sparrow.

**Figure 5:**
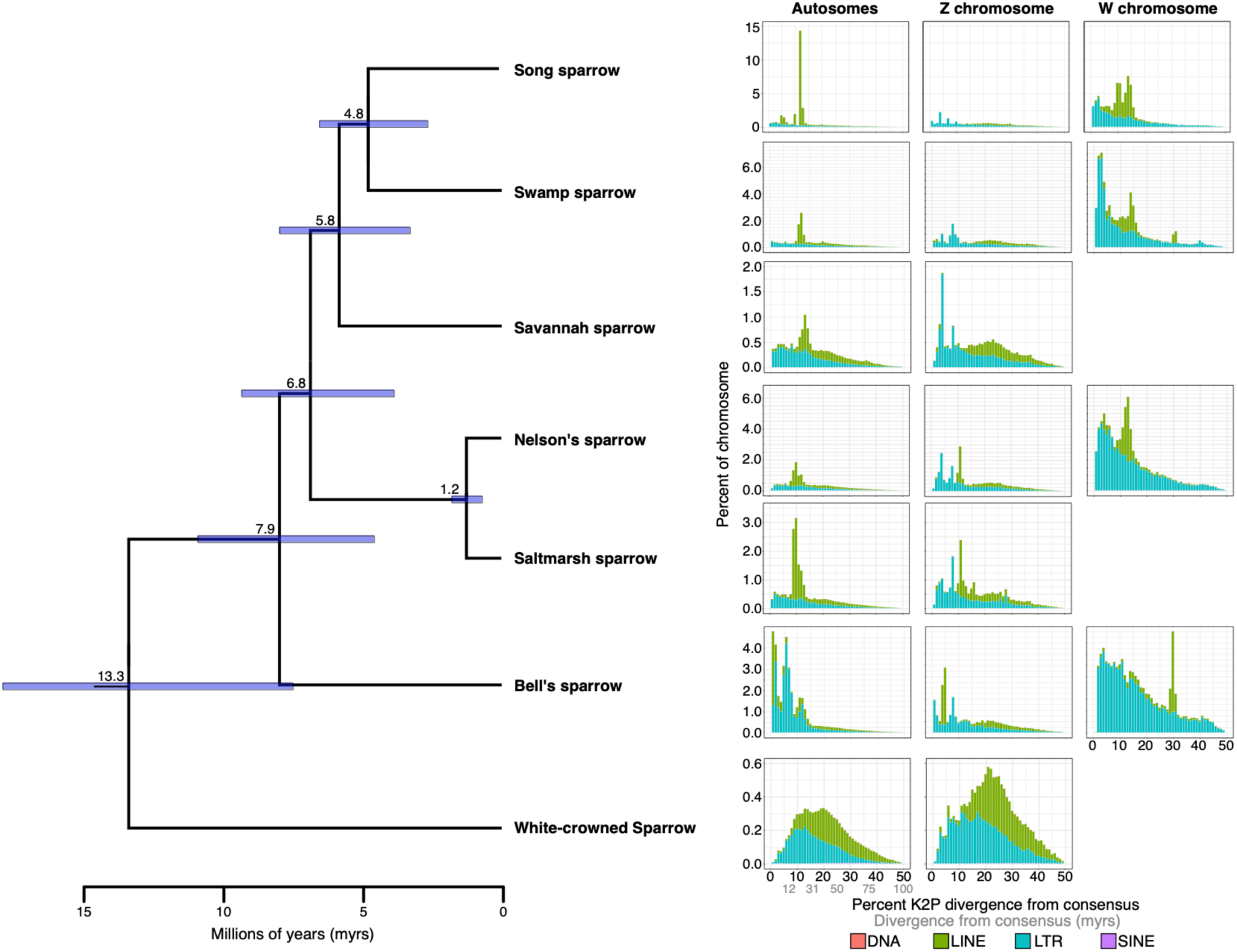
Transposable element (TE) landscapes for the autosomal, Z, and W chromosomes. Left panel shows time-calibrated UCE phylogeny of seven sparrow species. Branch labels indicate mean estimate of divergence time for each node with purple bars indicating 95% HPD error around that estimate. All nodes in the topology received bootstrap support of 100%. Right panel shows TE landscapes for each species. Percent divergence on the x-axis was calculated as the percent Kimura 2-parameter (K2P) distance with CpG sites excluded. The abundance of TEs in each percent divergence bin was normalized as a percentage of the chromosome length on the y-axis.

## DISCUSSION

### Highly contiguous and complete genomes reveal high repeat content

We generated highly contiguous and complete genomes of three sparrow species in the family Passerellidae that we compared with three chromosome-level genomes generated by the Vertebrate Genome Project. Contig N50 for the three newly generated genomes exceeded 92%, and the scaffold N50 exceeded 85% of all avian genomes recently surveyed by Bravo et al. (2021). Assembly length for the six species analyzed here also exceeds assembly lengths for all short-read based assemblies generated to date (1.16-1.40Gb vs. 0.91-1.05Gb). The longer length of these assemblies more closely approximates independent estimates of genome size from Feulgen image analysis densitometry (C-value), with song and Savannah sparrow missing only 2-3% of genomic sequence relative to C-value size estimates. Longer assemblies were also associated with greater levels of repeat content. The high percentage of total interspersed repeats discovered in the song sparrow (31.2%), Bell’s sparrow (29.5%), and California towhee (30.9%; also see Black et al. 2023) genomes are the highest levels ever reported for Passeriformes and more closely resemble levels of repeat content described in the avian orders Piciformes and Bucerotiformes (Manthey et al. 2018; Feng et al. 2020). Although our finding is a novel result for passerine genome assemblies, reassociation kinetic studies found about 36% of the dark-eyed junco genome to be repetitive DNA (Shields & Straus 1975). Recent long-read assemblies for jays in the passerine family Corvidae also show repeat content in-line with the results presented here (Benham et al. 2023; DeRaad et al. 2023). Further, high levels of repeat content in these sparrows matches predictions that much of the missing genomic data from avian short-read assemblies are likely repetitive DNA (Elliot & Gregory 2015; Kapusta & Suh 2017; Peona et al. 2018). We expect that the generation of additional highly contiguous and complete genomes using third generation sequencing technology will also find higher levels of repeat content in avian genomes than previously appreciated.

Previous sparrow genome assemblies generated using short-read methods were found to be missing ∼12-30% of DNA sequence relative to C-values (Fig. 2b). The majority of this missing DNA is likely associated with highly repetitive regions of the genome that caused gaps in prior assemblies. Gaps associated with repeat regions is a well-established phenomenon and recent comparisons among sequencing technologies point to long contiguous reads as essential for spanning these gaps (Rhie et al. 2021). Similarly, we find that sparrow assemblies generated using PacBio long-read sequence data exhibited elevated contig N50 and higher percent repeat content, and that percent repeat content decreased in assemblies inferred to be missing a greater percentage of DNA (Fig. 3). This was also true for intra-specific comparisons of multiple song and saltmarsh sparrow genomes where repeat content increased with assembly length and contiguity. The diversity of transposable elements can vary significantly within and among populations (e.g. *Ficedula* flycatchers; Suh et al. 2018). Although we compared song sparrow genomes from three different subspecies that could differ in repeat content and genome size, the patterns for song sparrow are consistent with work showing increased TE discovery and abundance in long-read versus short-read assemblies of the same individual (Peona et al. 2021). Our results are also consistent with prior work linking metrics of genome contiguity with levels of repeat content detected in genome assemblies (Galbraith et al. 2021). Overall, our analyses among closely related sparrow species confirms that sequencing technology appears to be a major confounding factor in comparative research on TE diversity.

### Genome size evolution in birds

Transposable elements are widely recognized as an important driver of genome size evolution across Eukaryotes (Kidwell 2002; Elliott & Gregory 2015). Low repeat content and high synteny contributed to an early view that avian genomes were relatively stable and constrained in size as an adaptation for the metabolic demands of flight (Hughes & Hughes 1995; Wright et al. 2014). This perspective has been challenged with evidence of a more dynamic history of avian genome expansions followed by large-scale deletions (Kapusta et al. 2017). Genome size variation in the Passerellidae, as measured by densitometry and cytometry methods, ranges from 1.13-1.63 Gb, with the Savannah sparrow possessing the smallest genome of any sparrow measured to date. Differences in assembly length across all sparrow species were entirely related to repeat content (Fig. 3). Further, TE composition differed significantly even in species with similar genome assembly lengths (e.g. Bell’s and song sparrow).

Unlike the other sparrow species analyzed, CR1 LINE elements were found to be the most abundant TE class within the song sparrow genome. The TE landscape of the song sparrow genome indicates that the majority of LINE DNA stems from a period of increased activity 25-30 million years ago. All six sparrow species in the grassland clade show a spike in LINE activity during this period, but much of the LINE DNA from this period was eliminated in species other than the song sparrow. In contrast, the white-crowned sparrow shows a more ‘bell-curve’ shape of LINE element proliferation with a peak of less than 0.5% at ∼30-40 Mya (about 20% divergence from consensus). The white-crowned sparrow pattern more closely resembles TE landscapes observed in other Passerifomes birds such as *Ficedula* flycatchers (Muscicapidae; Suh et al. 2017) and Estrildidae finches (Boman et al. 2019). These patterns point to a proliferation of LINE elements within the grassland sparrow clade that more likely occurred after divergence from the white-crowned sparrow ∼13.3 Mya. This discrepancy in the timing of activity could reflect the use of a genome wide estimate of mutation rate from *Ficedula* flycatchers (Smeds et al. 2016) that may be underestimating the true mutation rate for Passerellidae sparrows and/or transposable elements. Indeed, many of the host genome’s defense mechanisms against TE proliferation involve DNA-editing enzymes, such as APOBECs, which mutate TE sequences to silence their activity in the genome (Goodier et al. 2016; Knisbacher & Levanon 2016).

LTR elements were the most abundant TE within all sparrow genomes except the song sparrow. Proliferation of these elements has also been more recent, beginning ∼12 million years ago and continuing to the present. Recent proliferation of LTR elements was especially pronounced in the Bell’s sparrow. Recent proliferation of LTR elements more closely aligns with patterns of TE expansion observed in the zebra finch (Kapusta & Suh 2017) and the blackcap (*Sylvia atricapilla*; Bours et al. 2023). The reasons for recent LTR expansions in songbirds versus other avian lineages (e.g. Chicken; Warren et al. 2017) are not entirely clear. One possibility is that competition for similar genomic insertion sites between LTR and LINE elements could be mediated by host defenses. Recent work in the deer mouse (*Peromyscus maniculatus;* Gozashti et al. 2023) provided evidence for a cycle initiated by greater host repression of ancient endogenous retroviruses (ERV, a type of LTR) that allowed for greater LINE proliferation in the genome. This was followed by the invasion of the deer mouse genome by a novel ERV that was hypothesized to have a greater immunity to host defense mechanisms and greater potential to outcompete LINE elements for insertion sites. A related possibility could reflect the accumulation of LTR elements on the W chromosome, many of which remain transcriptionally active and could seed invasions of the autosomal chromosomes (see below; Peona et al. 2021). Whether either of these scenarios contributes to the recent expansions of LTR elements in songbird genomes awaits further study; however, the expanding number of avian genomes assembled using long-read sequence data will be essential for understanding the dynamics of TE proliferation and deletion in the evolution of avian genomes.

Differences in genome size and repeat content could be the result of a number of different mechanisms involved in TE silencing, deletion, or expansion in host genomes (Goodier et al. 2016). A wide range of epigenetic mechanisms exist to silence TE activity in plants and animals (reviewed in Slotkin & Martienssen 2007). In birds, methylation of CpG and non-CpG sites in TEs with DNA methyltransferases is the primary mechanism of TE silencing that has been documented (Derks et al. 2016; Kapusta & Suh 2017). Mutating TE sequences is another mechanism hosts deploy to defend against TE proliferation. APOBEC genes induce C-to-U mutations in retrotransposons leading to inactivation and degradation of these elements. The genomes of zebra finch and other bird species exhibit signatures of high APOBEC activity (Knisbacher & Levanon 2015). An important mechanism for the removal of LTR elements from the genome is ectopic recombination. This process deletes most of the element sequence leaving only a single LTR and correlates with recombination rate variation across the genome in birds (Ji & DeWoody 2016). Finally, demographic differences among populations could influence TE dynamics, with TEs predicted to insert and spread more rapidly in populations with a small effective population size (Ne; Lynch & Conery 2003). Demographic analyses of the different sparrow species provide some support for this hypothesis as both Nelson’s and saltmarsh sparrows have been inferred to experience historical bottlenecks and lower Ne than other species (Walsh et al. 2019a,b; Walsh et al. 2021). In contrast, the Savannah sparrow has the lowest repeat content and has been inferred to maintain high and constant effective population sizes (Benham & Cheviron 2019), which may be important for combating TE proliferation and maintaining a smaller genome. However, the relationship between Ne and TE proliferation is not necessarily straightforward (Whitney & Garland 2010) and some authors argue that TE expansions may be even more likely in species with large Ne (Ågren & Wright 2011).

Disruption of a host’s TE repression mechanisms can lead to TE expansions. Stressful conditions (e.g. thermal stress) can disrupt epigenetic silencing of TEs in the host genome, leading to TE expansions (Capy et al. 2000, Slotkin & Martienssen 2007). Furthermore, co-evolutionary arms races between TEs and the host genomes could lead to divergence in TE repressors among populations or closely related species. Subsequent hybridization among these lineages could allow TEs to escape their repressors and proliferate throughout the genome of hybrids (Bingham et al. 1982; Serrato-Capuchina & Matute 2018). Examples of TE re-activation following hybridization have been documented in both plants (Josefsson et al. 2006) and animals (O’Neill et al. 1998). In hybrid *Helianthus* sunflower species, proliferation of LTR elements was found to contribute significantly to a 50% increase in the genome size of hybrids relative to parental species (Ungerer et al. 2006). Intriguingly, the Bell’s sparrow individual used to generate the reference assembly for this project comes from the same subspecies known to hybridize with sagebrush sparrow in a contact zone centered ca. 120-150 km. to the northwest of the collecting locality. Whether recent hybridization between Bell’s and sagebrush or Nelson’s and saltmarsh sparrow lineages led to a TE expansion remains to be determined. However, the dynamic patterns of genome size evolution within sparrows indicates that the Passerellidae are an exciting model for future research on the dynamics of TE evolution.

### TE element proliferation on sex chromosomes

The potential deleterious effects of TE insertions is thought to explain a general trend of TE prevalence in regions of lower recombination rate (Rizzon et al. 2002; Ji & DeWoody 2016; Kent et al. 2017). These patterns are especially pronounced on the Y/W sex chromosomes where a lack of recombination, low gene density, and small effective population sizes are thought to allow for TE accumulation (Charlesworth & Langly 1988; Bachtrog 2003). This high TE abundance is thought to be a major contributing factor to the challenges of sequencing and assembling the W chromosome in birds and Y chromosome in mammals (Tomaskiewicz et al. 2017). Consistent with these expectations for the non-recombining W chromosome, we found that repeat content on the W chromosome was dramatically higher across all four female birds sequenced relative to autosomal or Z chromosomes. Interspersed repeats comprised 79.2% and 82.6% of the Nelson’s and Bell’s sparrow W chromosome, respectively, while the song and swamp sparrow (both *Melospiza*) possessed W chromosomes with over 90% repeat content. Previous reports of repeat content on the W chromosome range from 22% in the emu (*Dromaius novaehollandiae*; Peona et al. 2021) to over 84% in the hooded crow (e.g *Corvus cornix*; Warmuth et al. 2022) and 89% in the Steller’s jay (e.g. *Cyanocitta stelleri;* Benham et al. 2023). Similar to other avian W chromosome assemblies, endogenous retroviral elements are the dominant element representing 42.3% of the song to 69.2% of the Bell’s sparrow W chromosome assembly. Peona et al. (2021) also showed that a disproportionately large percentage of LTR elements on the avian W chromosome are full length retroviral elements that continue to be actively transcribed. The capacity of active elements to spread from the W to other regions of the genome makes the W chromosome a likely source for the recent activity and abundance of endogenous retroviral elements in Passeriformes (Warren et al. 2010; Zhang et al. 2014; Warmuth et al. 2022). Kapusta & Suh (2017) posited that the abundance of these elements in Passeriformes may have played critical roles in their high levels of diversification. The highly complete genome assemblies generated via third generation sequencing techniques will provide new opportunities to test this hypothesis.

## Conclusions

Here we report on the release of three highly contiguous assemblies of sparrows in the family Passerellidae. The combination of long-read and Omni-C technology enabled the generation of nearly complete and highly contiguous assemblies. Analysis of these genomes revealed a previously underappreciated abundance of repetitive elements in the genomes of songbirds and suggests that much of the missing data from other avian assemblies are likely comprised of repeat content. As third generation sequencing technologies become the standard in avian genome assembly, the dynamics of TE element proliferation and genome size evolution across different evolutionary timescales will become better understood. Our results point to the strong role repetitive element proliferation and deletion plays in the dynamics of avian genome size evolution, even among closely related species.

## ACKNOWLEDGEMENTS

We thank Joshua Ho for assistance with tissue subsampling, and Nicole Guido, Jonathan Clark and Chris Elphick for sample collection. PacBio Sequel II library prep and sequencing was carried out at the DNA Technologies and Expression Analysis Cores at the UC Davis Genome Center, supported by NIH Shared Instrumentation Grant 1S10OD010786-01. Deep sequencing of Omni-C libraries used the Novaseq S4 sequencing platforms at the Vincent J. Coates Genomics Sequencing Laboratory at UC Berkeley, supported by NIH S10 OD018174 Instrumentation Grant. We thank the staff at the UC Davis DNA Technologies and Expression Analysis Cores and the UC Santa Cruz Paleogenomics Laboratory for their diligence and dedication to generating high quality sequence data. We thank Erich Jarvis and personnel at the Vertebrate Genomes Project for conducting the sequencing and assembly of the Nelson’s, saltmarsh, and swamp sparrow.

## DATA AVAILIBILITY

Data generated for this study are available under NCBI BioProject PRJNA720569. Raw sequencing data for the Savannah sparrow sample FMNH:Bird:499929 (NCBI BioSample SAMN24839580) are deposited in the NCBI Short Read Archive (SRA) under SRS12336030. Raw sequencing data for the song sparrow sample MVZ:Bird:193390 (NCBI BioSample SAMN24817870, SAMN24817871) are deposited in the NCBI SRA under SRS12452128. Raw sequencing data for the Bell’s sparrow sample MVZ:Bird:192114 (NCBI BioSample SAMN24224802) are deposited in the NCBI SRA under SRS11988259. See supplemental Table S3 for the GenBank accession, BioProject, and BioSample numbers associated with the VGP and other genomes analyzed. For the CCGP genomes, assembly scripts and other data for the analyses presented can be found at the following GitHub repository: www.github.com/ccgproject/ccgp_assembly. Transposable element library, UCE sequences, and other supplemental data and code can be found on Dryad: https://doi.org/10.5061/dryad.cjsxksncs

