## Supplemental materials for "Remarkably high repeat content in the genomes of sparrows: the importance of genome assembly completeness for transposable element discovery"

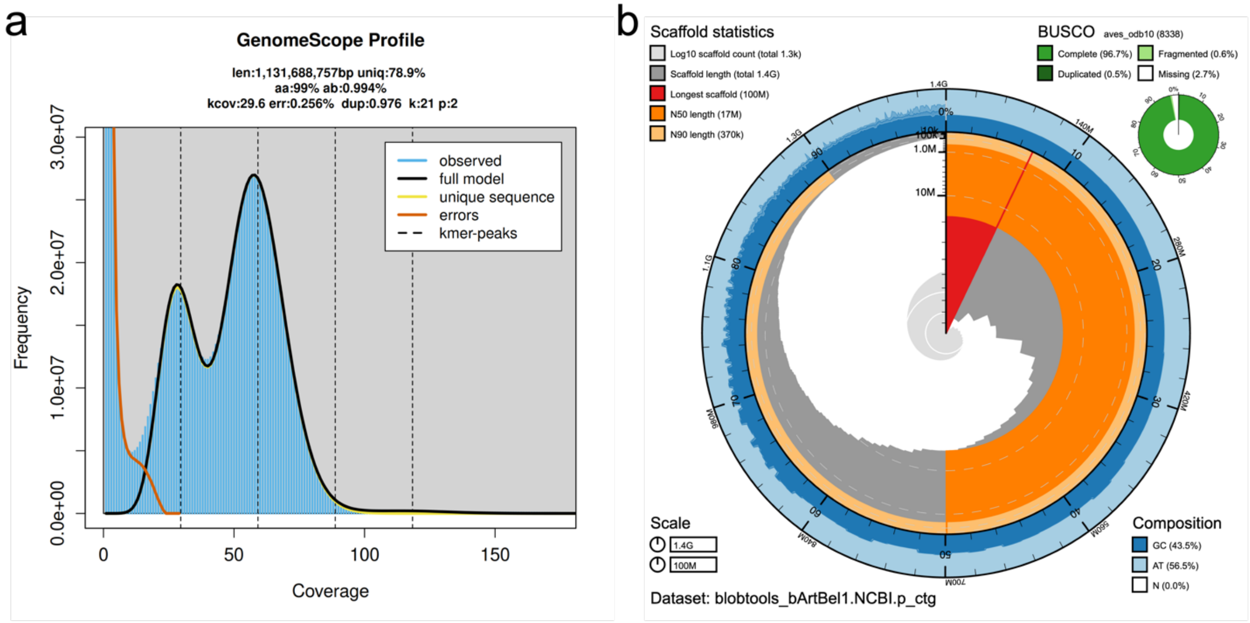


**Figure S1:** Visual overview of genome assembly metrics for the primary assembly of the Bell's sparrow (*Artemisiospiza belli*) genome. (a) K-mer spectra output generated from PacBio HiFi data without adapters using GenomeScope2.0. The bimodal pattern observed corresponds to a diploid genome and the k-mer profile matches that of low (<1%) heterozygosity. The left-hand K-mer peak at lower coverage and frequency corresponds to differences between haplotypes (heterozygous sites), whereas the right-hand k-mer peak at higher coverage and frequency correspond to similarities between haplotypes (homozygous sites). (b) BlobToolKit Snail plot showing a graphical representation of the quality metrics presented in Table 1 for the *Artemisiospiza belli* primary assembly (bArtBel1). The plot circle represents the full size of the assembly. From the inside-out, the central plot covers length-related metrics. The red line represents the size of the longest scaffold; all other scaffolds are arranged in size-order moving clockwise around the plot and drawn in gray starting from the outside of the central plot. Dark and light orange arcs show the scaffold N50 and scaffold N90 values. The central light gray spiral shows the cumulative scaffold count with a white line at each order of magnitude. White regions in this area reflect the proportion of Ns in the assembly; the dark versus light blue area around it shows mean, maximum, and minimum GC vs. AT content at 0.1% intervals (Challis et al. 2020).

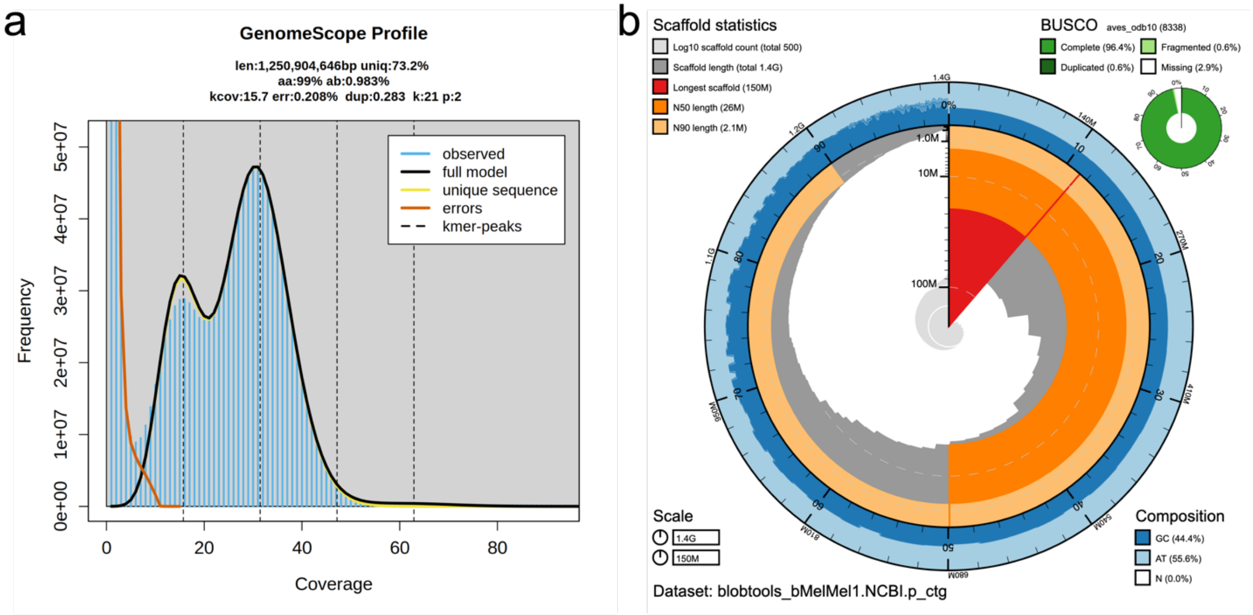


**Figure S2:** Visual overview of genome assembly metrics for the primary assembly of the song sparrow (*Melospiza melodia*) genome. (a) K-mer spectra output generated from PacBio HiFi data without adapters using GenomeScope2.0. The bimodal pattern observed corresponds to a diploid genome and the k-mer profile matches that of low (<1%) heterozygosity. The left-hand K-mer peak at lower coverage and frequency corresponds to differences between haplotypes (heterozygous sites), whereas the right-hand k-mer peak at higher coverage and frequency correspond to similarities between haplotypes (homozygous sites). (b) BlobToolKit Snail plot showing a graphical representation of the quality metrics presented in Table 1 for the *Melospiza melodia* primary assembly (bMelMel1). The plot circle represents the full size of the assembly. From the inside-out, the central plot covers length-related metrics. The red line represents the size of the longest scaffold; all other scaffolds are arranged in size-order moving clockwise around the plot and drawn in gray starting from the outside of the central plot. Dark and light orange arcs show the scaffold N50 and scaffold N90 values. The central light gray spiral shows the cumulative scaffold count with a white line at each order of magnitude. White regions in this area reflect the proportion of Ns in the assembly; the dark versus light blue area around it shows mean, maximum, and minimum GC vs. AT content at 0.1% intervals (Challis et al. 2020).

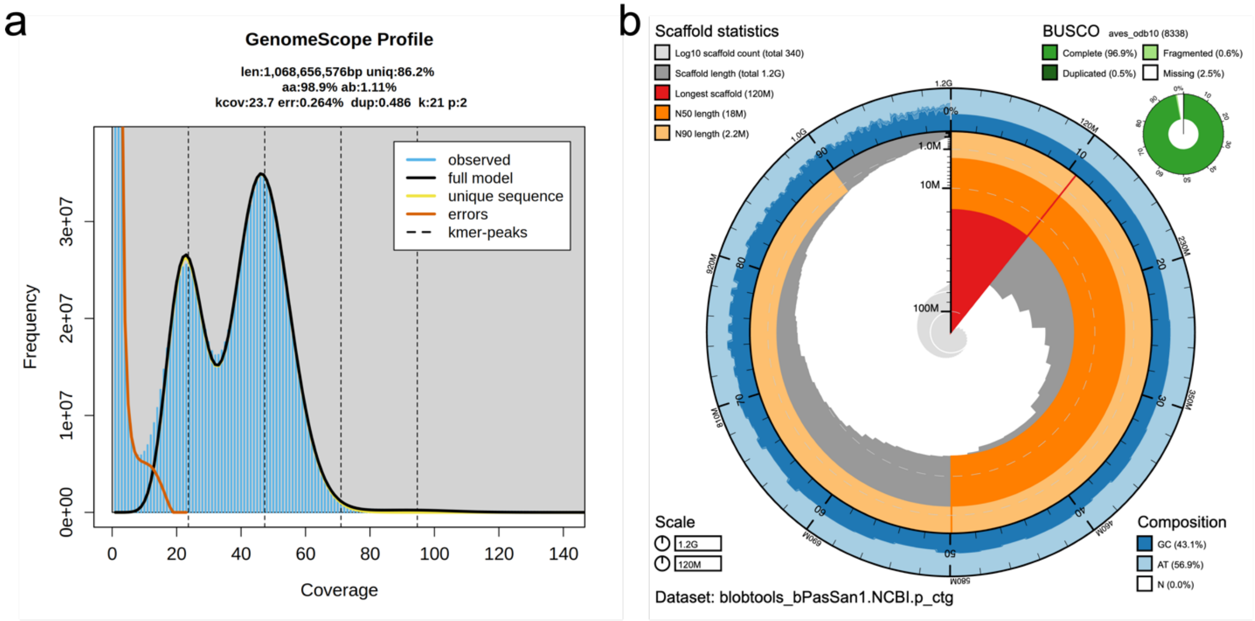


**Figure S3:** Visual overview of genome assembly metrics for the primary assembly of the Savannah sparrow (*Passerculus sandwichensis*) genome. (a) K-mer spectra output generated from PacBio HiFi data without adapters using GenomeScope2.0. The bimodal pattern observed corresponds to a diploid genome and the k-mer profile matches that of low (<1%) heterozygosity. The left-hand K-mer peak at lower coverage and frequency corresponds to differences between haplotypes (heterozygous sites), whereas the right-hand k-mer peak at higher coverage and frequency correspond to similarities between haplotypes (homozygous sites). (b) BlobToolKit Snail plot showing a graphical representation of the quality metrics presented in Table 1 for the *Passerculus sandwichensis* primary assembly (bPasSan1). The plot circle represents the full size of the assembly. From the inside-out, the central plot covers length-related metrics. The red line represents the size of the longest scaffold; all other scaffolds are arranged in size-order moving clockwise around the plot and drawn in gray starting from the outside of the central plot. Dark and light orange arcs show the scaffold N50 and scaffold N90 values. The central light gray spiral shows the cumulative scaffold count with a white line at each order of magnitude. White regions in this area reflect the proportion of Ns in the assembly; the dark versus light blue area around it shows mean, maximum, and minimum GC vs. AT content at 0.1% intervals (Challis et al. 2020).

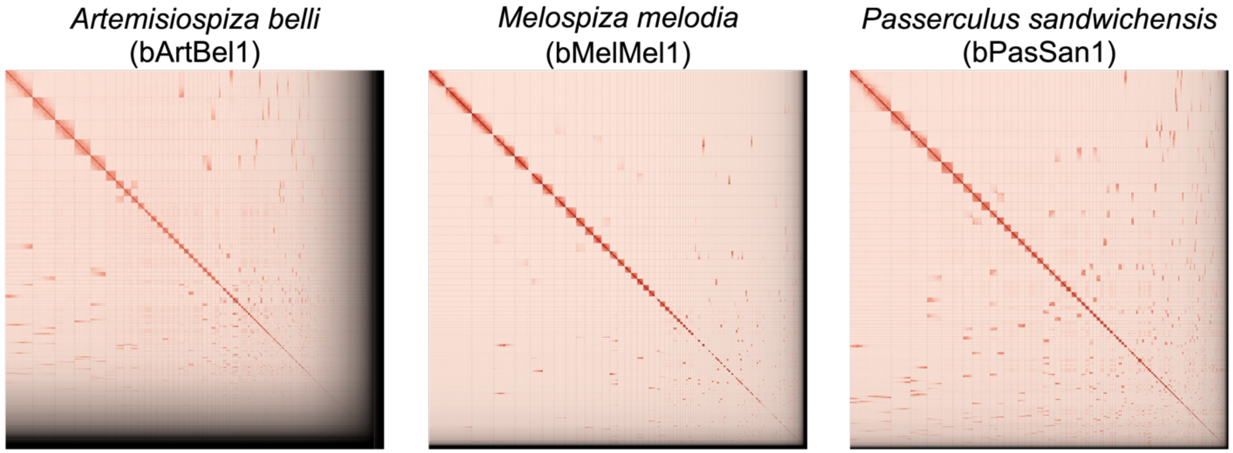


**Figure S4:** Hi-C Contact maps for the primary assemblies of all three sparrow species. Hi-C contact maps were generated with PretextSnapshot and translate proximity of genomic regions in 3D space to contiguous linear organization. Each cell in the contact map corresponds to sequencing data supporting the linkage (or join) between two such regions.

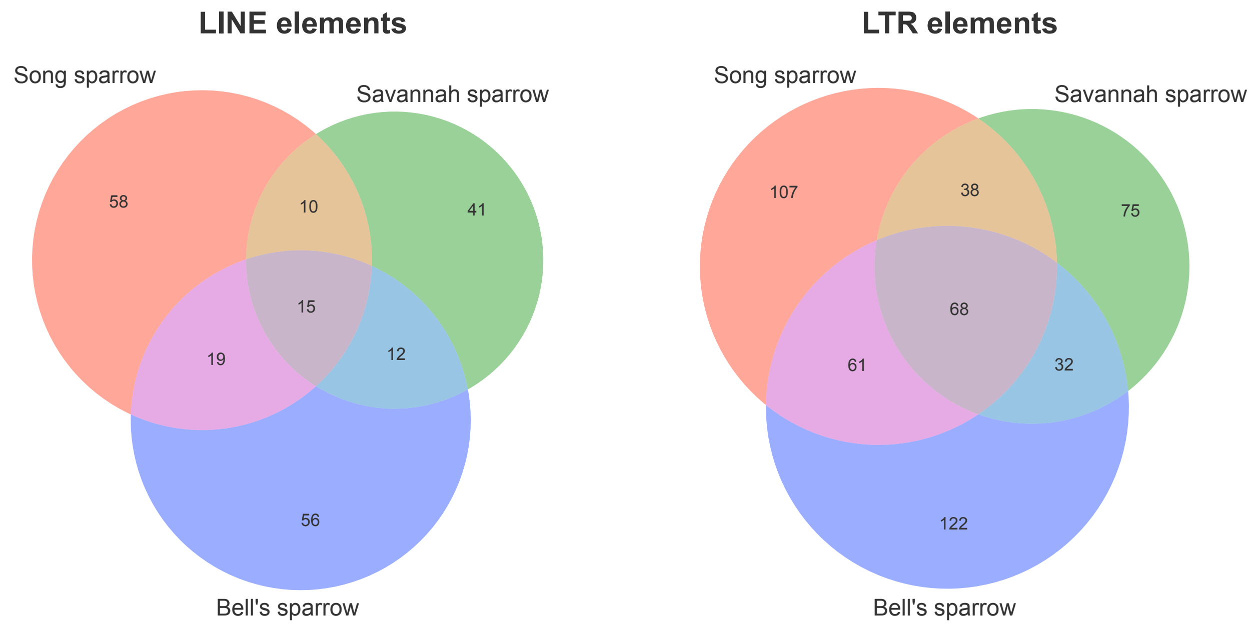


**Figure S5:** Venn diagram showing overlapping LINE and LTR element families within the sparrow TE library identified with RepeatModeler2 and manual curation.

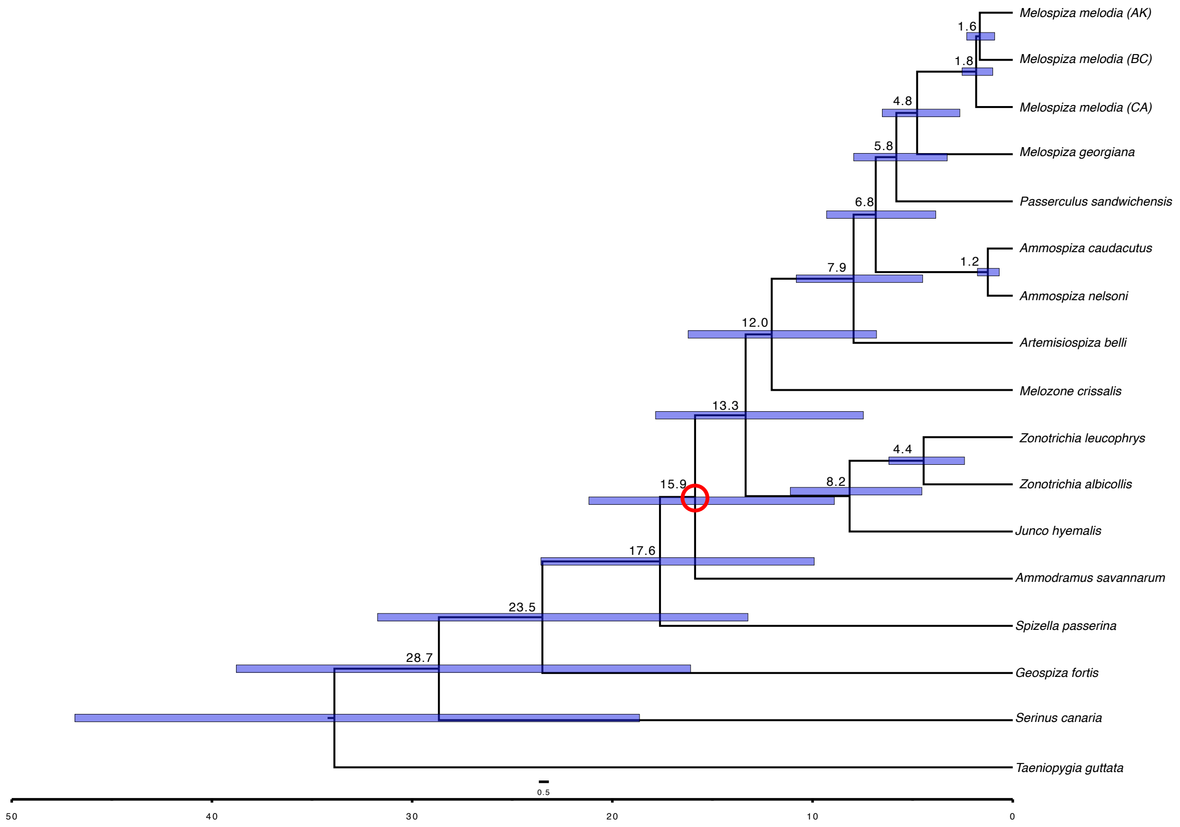


**Figure S6:** Time-calibrated phylogeny of sparrows with outgroups included. Branch annotations signal point estimate of divergence time for each node. Purple bars show 95% HPD of divergence time estimate for each node. Red circle denotes node with fossil calibration. Input topology to MCMCtree was generated in RAxML and had bootstrap support of 100 for each node.

**Table S1:** Summary of sequencing data used to generate assemblies for the Savannah, Bell's and song sparrow genomes assembled by the California Conservation Genomics Project.

| Species | | *Artemisiospiza belli* | *Melospiza melodia* | *Passerculus sandwichensis* |
|---|---|---|---|---|
| **SRA Accessions** | | | | |
| PacBio HiFi reads | Run information | 1 PACBIO_SMRT (Sequel II) run: 4M spots, 70.8G bases, 54.3Gb | 1 PACBIO_SMRT (Sequel II) run: 2.5M spots, 41.1G bases, 31Gb | 1 PACBIO_SMRT (Sequel II) run: 3.2M spots, 53.5G bases, 40.6Gb |
| | Accession | SRX14164085 | SRX14688640 | SRX14558718 |
| Omni-C Illumina reads | Run information | 2 ILLUMINA (Illumina NovaSeq 6000) runs: 197.7M spots, 59.7G bases, 19Gb | 2 ILLUMINA (Illumina NovaSeq 6000) runs: 138.9M spots, 41.9G bases, 13.5Gb | 2 ILLUMINA (Illumina NovaSeq 6000) runs: 158.3M spots, 47.8G bases, 15.2Gb |
| | Accession | SRX14164086, SRX14164087 | SRX14688641, SRX14688642 | SRX14558719,SRX14558720 |
| **PacBio HiFi sequencing data summary** | | | | |
| PacBio HiFi Number of reads | | 4,032,791 | 2,513,457 | 3,225,671 |
| PacBio coverage | | 50 | 32 | 62 |
| Read N50 | | 16,833 | 16,549 | 17,788 |
| Minmum size | | 44 | 43 | 43 |
| Mean size | | 16,596 | 16,370 | 17,546 |
| Maximum size | | 53,098 | 56,971 | 56,639 |
| **GenomeScope (PacBio HiFi based)** | | | | |
| Genome size estimation (bp) | | 1,131,688,757 | 1,250,904,646 | 1,068,656,576 |
| Genome Repeat length (bp) | | 239,245,991 | 335,139,019 | 147,539,127 |
| Genome Unique length (bp) | | 892,442,766 | 915,765,628 | 921,117,449 |
| Sequencing error rate | | 0.26% | 0.21% | 0.26% |
| Heterozygosity | | 0.99% | 0.98% | 1.11% |

§ Read coverage (per species) has been calculated based on the GenomeScope estimated genome size

**Table S2:** Assembly pipeline and software used for the Savannah, Bell's and song sparrow genomes assembled by the California Conservation Genomics Project.

| Assembly | Software and options § | Version |
|---|---|---|
| Filtering PacBio HiFi adapters | HiFiAdapterFilt | Commit 64d1c7b |
| Kmer counting | Meryl (k=21) | 1 |
| Estimation of genome size and heterozygosity | GenomeScope | 2 |
| *De novo assembly (contiging)* | HiFiasm (--primary, HiC mode, output p_ctg,a_ctg) | 0.16.1-r375 |
| Remove low-coverage, duplicated contigs | purge_dups | 1.2.6 |
| **Scaffolding** | | |
| Omni-C data alignment | Arima Genomics Mapping Pipeline | Commit 2e74ea4 |
| Omni-C Scaffolding | SALSA (-DNASE, -i 20, -p yes) | 2 |
| Gap closing | YAGCloser (-mins 2 -f 20 -mcc 2 -prt 0.25 -eft 0.2 -pld 0.2) | Commit 20e2769 |
| Long-read alignment | minima2 (-ax map-pb, secondary=yes) | 2.18-r1015 |
| **Omni-C Contact map generation** | | |
| Short-read alignment | BWA-MEM (-5SP) | 0.7.17-r1188 |
| SAM/BAM processing | samtools | 1.11 |
| SAM/BAM filtering | pairtools | 0.3.0 |
| Pairs indexing | pairix | 0.3.7 |
| Matrix generation | cooler | 0.8.10 |
| Matrix balancing | HicExplorer (hicCorrectmatrix correct --filterThreshold -2 4) | 3.6 |
| | HiGlass | 2.1.11 |
| | PretextMap | 0.1.4 |
| | PretextView | 0.1.5 |
| Contact map visualization | PretextSnapshot | 0.03 |
| **Organelle assembly** | | |
| Mitogenome assembly | MitoHiFi (-r, -p 50, -o 1) | Commit c06ed3e |
| **Genome quality assessment** | | |
| Basic assembly metrics | QUAST (--est-ref-size) | 5.0.2 |
| | BUSCO (-m geno, -l aves) | 5.0.0 |
| Assembly completeness | Merqury | 2022-01-29 |
| Synteny visualization | JupiterPlot (ng=80, m=1000000) | Commit CCCCC |
| **Contamination screening** | | |
| Local alignment tool | BLAST+ (blastn -db nt, -outfmt '6 qseqid staxids bitscore std' , -max_target_seqs 1, -max_hsps 1, -evalue 1e-25 ) | 2.10.0 |
| General contamination screening | BlobToolKit | 2.3.3 |

| | | |
|---|---|---|
| **Repeat analyses** | | |
| | RepeatModeler (ltrstruct) | 2 |
| | RepeatMasker | 4.1.2 |
| | cd-hit-est | 4.8.1 |
| | mafft | 7.49 |
| | EMBOSS (cons) | |
| Identification of open reading frames | TE-Aid | |
| | minimap2 | 2 |
| **Phylogeny construction** | | |
| | Raxml | 8 |
| | PAML (MCMCtree) | 4.9 |
| | tracer | 1.6.0 |

§Options detailed for non-default parameters.

**Table S3:** Summary of assembly name, accession, sequencing methods, and assembly approaches for each of the 15 genomes analyzed in this study.

| Species | scientific name | Assembly name | Genbank accession | BioProject | SAMN | Tissue | sequencing technology | Assembly methods |
|---|---|---|---|---|---|---|---|---|
| **Savannah sparrow** | *Passerculus sandwichensis* | bPasSan1.0.p | GCA_022577445.1 | PRJNA796788 | SAMN24839580 | liver | PacBio Sequel II; PacBio Sequel IIe; Dovetail OmniC; Illumina NovaSeq | HiFiasm v. 0.16.1-r375; purge_dups v. 1.2.5; SALSA2 v. 2 |
| **Song sparrow** | *Melospiza melodia gouldii* | bMelMel1.0.p | GCA_022749695.1 | PRJNA796324 | SAMN24817870 | blood | PacBio Sequel II; PacBio Sequel IIe; Dovetail OmniC; Illumina NovaSeq | HiFiasm v. 0.16.1-r375; purge_dups v. 1.2.5; SALSA2 v. 2 |
| **Bell's sparrow** | *Artemisiospiza belli* | bArtBel1.0.p | GCA_021963965.1 | PRJNA791509 | SAMN24224802 | liver | PacBio Sequel II; PacBio Sequel IIe; Dovetail OmniC; Illumina NovaSeq | HiFiasm v. 0.16.1-r375; purge_dups v. 1.2.5; SALSA2 v. 2 |
| **Nelson's sparrow** | *Ammospiza nelsoni* | bAmmNel1.pri | GCA_027579445.1 | PRJNA839456 | SAMN28421656 | blood | PacBio Sequel II HiFi; Bionano Genomics DLS; Arima Hi-C v2 | HiFiasm v. 0.15.4 + galaxy0; purge_dups v. 1.2.5 + galaxy3; Bionano Solve v. 3.6.1 + galaxy3; salsa v. 2.3 + galaxy2 |
| **Saltmarsh sparrow** | *Ammospiza caudacuta* | bAmmCau1.pri | GCA_027887145.1 | PRJNA839452 | SAMN28421630 | blood | PacBio Sequel II HiFi; Bionano Genomics DLS; Arima Hi-C v2 | HiFiasm v. 0.15.4 + galaxy0; purge_dups v. 1.2.5 + galaxy3; Bionano Solve |

| | | | | | | | | |
|---|---|---|---|---|---|---|---|---|
| | | | | | | | | v. 3.6.1 + galaxy3; salsa v. 2.3 + galaxy2 |
| **Swamp sparrow** | *Melospiza georgiana* | bMelGeo1.pri | GCA_028018845.1 | PRJNA915609 | SAMN22787412 | blood | PacBio Sequel II HiFi; Bionano Genomics DLS; Arima Hi-C v2 | HiFiasm v. 0.15.4 + galaxy0; purge_dups v. 1.2.5 + galaxy3; Bionano Solve v. 3.6.1 + galaxy3; salsa v. 2.3 + galaxy2 |
| **Song sparrow (BC)** | *Melospiza melodia rufina* | ASM1339820v1 | GCA_013398205.1 | PRJNA545868 | SAMN12253982 | blood | Illumina; Pacbio_SMRT | Platanus; PBJelly v. 1.2.4; 15.8.24 |
| **Song sparrow (AK)** | *Melospiza melodia maxima* | Mmel_1.0 | GCA_011057915.1 | PRJNA511035 | SAMN10622322 | blood | Illumina HiSeq | HiRise v. OCT-2016 |
| **Saltmarsh sparrow (sr)** | *Ammospiza caudacuta* | NA | NA | NA | NA | blood | Illumina HiSeq | Allpaths-LG v. 44849 |
| **white-throated sparrow** | *Zonotrichia albicollis* | Zonotrichia_albicollis-1.0.1 | GCF_000385455.1 | PRJNA197293 | SAMN02981528 | blood | Illumina | Allpaths-LG v. Feb-2013 |
| **white-crowned sparrow** | *Zonotrichia leucophrys* | RI_Zleu_1.0 | GCA_028769735.1 | PRJNA889240 | SAMN31812169 | muscle | PacBio Sequel | HiRise v. 2016 |
| **dark-eyed junco** | *Junco hyemalis* | dark-eyed_junco_22Sep2020_assembly | GCA_003829775.2 | PRJNA493001 | SAMN10120167 | muscle | Illumina NovaSeq | HiRise v. JAN-2016 |
| **chipping sparrow** | *Spizella passerina* | ASM1340137v1 | GCA_013401375.1 | PRJNA545868 | SAMN12253929 | tissue | Illumina HiSeq | SOAPdenovo v. 2.04 |
| **grasshopper sparrow** | *Ammodramus savannarum* | ASM2046641v1 | GCA_020466415.1 | PRJNA747010 | SAMN20250500 | blood | Illumina HiSeq | HiRise v. JULY-2018 |
| **California towhee** | *Melozone crissalis* | PUWL_Pcris_2 | GCA_028551555.1 | PRJNA915193 | SAMN32379461 | blood | PacBio | HiFiasm v. 2022 |

**Table S4:** Table of assembly quality statistics and BUSCO search results from each of the draft CCGP sparrow assemblies. BUSCO results were performed using the 8,338 universal single copy genes in birds found in the aves_odb10 database.

| Species | *Artemisiospiza belli* | | *Melospiza melodia* | | *Passerculus sandwichensis* | |
|---|---|---|---|---|---|---|
| **Bio Projects & Vouchers** | | | | | | |
| CCGP NCBI BioProject | PRJNA720569 | | PRJNA720569 | | PRJNA720569 | |
| Genera NCBI BioProject | PRJNA766272 | | PRJNA765629 | | PRJNA765656 | |
| Species NCBI BioProject | PRJNA777142 | | PRJNA777193 | | PRJNA777205 | |
| NCBI BioSample | SAMN24224802 | | SAMN24817870, SAMN24817871 | | SAMN24839580 | |
| Specimen identification | MVZ:Bird:192114 | | MVZ:Bird:193390 | | FMNH 499929 | |
| NCBI Genome accessions | **Primary** | **Alternate** | **Primary** | **Alternate** | **Primary** | **Alternate** |
| Assembly accession | JAKDEW000000000 | JAKDEX000000000 | JALCYL000000000 | JALCYM000000000 | JAKOOL000000000 | JAKOOM00000000 |
| Genome sequences | GCA_021966175.1 | GCA_021963965.1 | GCA_022749695.1 | GCA_022749775.1 | GCA_022577445.1 | GCA_022578375.1 |
| Organelles | JAKDEW010001339.1 (partial) | | JALCYL010000501.1 (partial) | | JAKOOL010000337.1 (partial) | |
| **Genome assembly metrics** | | | | | | |
| Assembly identifier | bArtBel1 | | bMelMel1 | | bPasSan1 | |
| Quality code * | 7.7.Q58.C66 | | 6.7.Q60.C50 | | 6.7.Q60.C72 | |
| | **Primary** | **Alternate** | **Primary** | **Alternate** | **Primary** | **Alternate** |
| Number of contigs | 1,539 | 43,036 | 823 | 34,692 | 676 | 31,565 |
| Contig N50 (bp) | 8,253,817 | 107,859 | 8,311,625 | 141,745 | 5,981,027 | 133,443 |
| Contig NG50 § | 11,625,514 | 280,805 | 9,239,046 | 247,555 | 6,348,013 | 240,328 |
| Longest Contigs | 35,931,659 | 6,669,798 | 59,497,540 | 4,826,274 | 32,137,824 | 6,090,697 |
| Number of scaffolds | 1,339 | 43,024 | 501 | 34,687 | 337 | 29,743 |
| Scaffold N50 | 17,082,054 | 108,004 | 25,784,215 | 141,872 | 18,220,233 | 151,227 |
| Scaffold NG50 § | 21,696,301 | 280,950 | 28,374,017 | 247,555 | 19,083,476 | 626,611 |
| Largest scaffold | 99,814,828 | 6,669,798 | 153,992,920 | 4,826,274 | 124,432,526 | 10,046,320 |
| Size of final assembly | 1,401,818,823 | 2,257,281,364 | 1,356,304,709 | 1,938,101,713 | 1,152,292,115 | 1,785,384,694 |
| Gaps per Gbp (# Gaps) | 143 (200) | 5(12) | 238 (323) | 3 (5) | 273 (314) | 1.013 (1,808) |
| Indel QV (Frame shift) | 41.67 | 41.95 | 41.00 | 41.38 | 41.67 | 41.68 |
| Base pair QV | 58.81 | 57.32 | 60.42 | 57.59 | 60.23 | 57.87 |

| | | | | | | | |
|---|---|---|---|---|---|---|---|
| | | Full assembly = 57.835 | | Full assembly = 58.55 | | Full assembly = 58.6511 | |
| k-mer completeness | | 89.46 | 85.49 | 90.32 | 84.27 | 85.05 | 86.95 |
| | | Full assembly = 99.273 | | Full assembly = 99.23 | | Full assembly = 99.5669 | |
| BUSCO completeness | Complete | 96.70% | 92.20% | 96.50% | 89.80% | 96.90% | 95.80% |
| | - Single | 96.20% | 82.10% | 95.90% | 81.60% | 96.40% | 85.10% |
| | - Duplicated | 0.50% | 10.10% | 0.60% | 8.20% | 0.50% | 10.70% |
| (aves_odb10) | Fragmented | 0.60% | 1.30% | 0.60% | 1.40% | 0.60% | 1.10% |
| n= 8,338 | Missing | 2.70% | 6.50% | 2.90% | 8.80% | 2.50% | 3.10% |

* Assembly quality code x.y.P.Q.C derived notation, from (Rhie et al. 2021). x = log10[contig NG50]; y = log10[scaffold NG50]; Q = Phred base accuracy QV (Quality value); C = % genome represented by the first 'n' scaffolds, following a known karyotype of 2n =74 for *P. sandwichensis* and estimated 2n=80 for both *M. melodia* and *A. belli*

§ NGx statistics have been calculated, per species, based on the GenomeScope estimates of genome sizes.

**Table S5:** Ultraconserved elements extracted from genomes of different sparrow and outgroup genomes. Table includes GenBank accession number for assembly used. Number of contigs extracted, min and max length of contigs, and total length of concatenated UCEs for each species.

| Common name | Scientific name | Accession number | UCE contigs | min length | max length | Total UCE length (bp) |
|---|---|---|---|---|---|---|
| Zebra finch | *Taeniopygia guttata* | GCA_003957565.4 | 4789 | 662 | 1778 | 5347378 |
| Island canary | *Serinus canaria* | GCA_022539315.2 | 4765 | 643 | 1781 | 5321849 |
| Medium ground finch | *Geospiza fortis* | GCA_000277835.1 | 4732 | 286 | 1781 | 5256921 |
| Chipping sparrow | *Spizella passerina* | GCA_013401375.1 | 3699 | 119 | 1698 | 3656630 |
| Grasshopper sparrow | *Ammodramus savannarum* | GCA_020466415.1 | 4419 | 600 | 1781 | 4925316 |
| White-throated sparrow | *Zonotrichia albicollis* | GCA_000385455.1 | 4781 | 634 | 1776 | 5321737 |
| White-crowned sparrow | *Zonotrichia leucophrys* | GCA_028769735.1 | 4730 | 1442 | 2755 | 10013019 |
| Dark-eyed junco | *Junco hyemalis* | GCA_003829775.2 | 4431 | 613 | 1781 | 4931968 |
| California towhee | *Melozone crissalis* | GCA_028551555.1 | 4768 | 1155 | 2760 | 10107862 |
| Bell's sparrow | *Artemisiospiza belli* | GCA_021963965.1 | 4839 | 686 | 1781 | 5412238 |
| Nelson's sparrow | *Ammospiza nelsoni* | GCA_027579445.1 | 4717 | 1356 | 2761 | 9982649 |
| Saltmarsh sparrow | *Ammospiza caudacuta* | GCA_027887145.1 | 4740 | 1117 | 2761 | 10031135 |
| Savannah sparrow | *Passerculus sandwichensis* | GCA_022577445.1 | 4822 | 623 | 1781 | 5390063 |
| Swamp sparrow | *Melospiza georgiana* | GCA_028018845.1 | 4719 | 1375 | 2761 | 9993216 |
| Song sparrow [CA] | *Melospiza melodia* | GCA_022749695.1 | 4811 | 636 | 1781 | 5381794 |
| Song sparrow [BC] | *Melospiza melodia* | GCA_013398205.2 | 4771 | 646 | 1781 | 5330005 |
| Song sparrow [AK] | *Melospiza melodia* | GCA_011057915.1 | 4749 | 465 | 1781 | 5250235 |
